# Calcium-induced calcium release and type 3 ryanodine receptors modulate the slow afterhyperpolarising current, sI_AHP_, and its potentiation in hippocampal pyramidal neurons

**DOI:** 10.1101/2020.03.03.974691

**Authors:** Angelo Tedoldi, Petra Ludwig, Gianluca Fulgenzi, Hiroshi Takeshima, Paola Pedarzani, Martin Stocker

**Author notes:** To whom correspondence should be addressed at: Research Department of Neuroscience, Physiology and Pharmacology University College London, Gower Street, London WC1E 6BT, United Kingdom (MS) (PP), or. Present address: Department of Physiology, University of Auckland, 85 Park Road, Grafton, Auckland, New Zealand (Angelo Tedoldi). Present address: Neural Development Section, Mouse Cancer Genetics Program, Center for Cancer Research, National Cancer Institute, National Institutes of Health, Frederick, United States (Gianluca Fulgenzi).

## Abstract

The slow afterhyperpolarising current, sI_AHP_, is a Ca^2+^-dependent current that plays an important role in the late phase of spike frequency adaptation. sI_AHP_ is activated by voltage-gated Ca^2+^ channels, while the contribution of calcium from ryanodine-sensitive intracellular stores, released by calcium-induced calcium release (CICR), is controversial in hippocampal pyramidal neurons. Three types of ryanodine receptors (RyR1-3) are expressed in the hippocampus, with RyR3 showing a predominant expression in CA1 neurons. We investigated the specific role of CICR, and particularly of its RyR3-mediated component, in the regulation of the sI_AHP_ amplitude and time course, and the activity-dependent potentiation of the sI_AHP_ in rat and mouse CA1 pyramidal neurons.

Here we report that enhancement of CICR by caffeine led to an increase in sI_AHP_ amplitude, while inhibition of CICR by ryanodine caused a small, but significant reduction of sI_AHP_. Inhibition of ryanodine-sensitive Ca^2+^ stores by ryanodine or depletion by the SERCA pump inhibitor cyclopiazonic acid caused a substantial attenuation in the sI_AHP_ activity-dependent potentiation in both rat and mouse CA1 pyramidal neurons. Neurons from mice lacking RyR3 receptors exhibited a sI_AHP_ with features undistinguishable from wild-type neurons, which was similarly reduced by ryanodine. However, the lack of RyR3 receptors led to a faster and reduced activity-dependent potentiation of sI_AHP_.

We conclude that ryanodine receptor-mediated CICR contributes both to the amplitude of the sI_AHP_ at steady state and its activity-dependent potentiation in rat and mouse hippocampal pyramidal neurons. In particular, we show that RyR3 receptors play an essential and specific role in shaping the activity-dependent potentiation of the sI_AHP_. The modulation of activity-dependent potentiation of sI_AHP_ by RyR3-mediated CICR contributes to plasticity of intrinsic neuronal excitability and is likely to play a critical role in higher cognitive functions, such as learning and memory.

## Introduction

The slow afterhyperpolarization (sAHP) has been first characterized nearly 40 years ago as a Ca^2+^-dependent K^+^ potential following action potentials or epileptiform bursts in hippocampal CA1 pyramidal neurons [1, 2]. Functionally, the sAHP is responsible for the late phase of spike frequency adaptation and leads to a strong reduction or a complete cessation of action potential firing, thereby controlling the repetitive firing of neurons and limiting the numbers of action potentials generated in response to stimuli [3, 4]. Voltage-clamp studies have revealed that the current, sI_AHP_, underlying the sAHP reaches its maximum with a time constant of several hundred milliseconds and decays with a time constant of >1s, and the kinetics of the current are temperature dependent [5, 6].

Activation of the sI_AHP_ requires Ca influx and an increase in intracellular Ca^2+^ concentration ([Ca^2+^]), as the current is suppressed by removing extracellular Ca^2+^ [1, 6], blocking Ca^2+^ channels [1, 2, 5, 6] or perfusing neurons with Ca^2+^ chelators, EGTA or BAPTA [6]. The Ca^2+^ sources that contribute to the activation of this current (sI_AHP_) and generate the afterhyperpolarising potential (sAHP) include voltage-gated calcium channels (VGCCs), whose subtypes vary in different neurons. In the hippocampus, the use of selective inhibitors for different VGCC subtypes has revealed that activation of L-type calcium channels substantially contributes to the generation of sI_AHP_/sAHP in both CA1 and CA3 pyramidal neurons [7–10]. Mice in which the gene encoding Ca_V_1.3 was deleted have further demonstrated that Ca_V_1.3 channels play a predominant role for the generation of sAHP in CA1 pyramidal neurons [11].

Two peculiar features of the sI_AHP_ and sAHP cannot be explained by a linear dependence on Ca^2+^ influx through VGCCs. The first is that the time to peak of their amplitude reaches its maximum value ∼500 ms after the end of Ca^2+^ entry during action potentials [12]. The second is the phenomenon of activity-dependent potentiation, often referred to as “run-up”, whereby repeated stimulation of cortical pyramidal neurons by depolarizing current pulses causes a marked and sustained increase in the sI_AHP_/sAHP amplitude with a concomitant reduction in neuronal excitability $ [13–17]. For each of these features Ca^2+^-induced Ca^2+^ release (CICR), where Ca^2+^ entering through VGCCs causes a secondary transient elevation of intracellular Ca^2+^ levels due to the activation of ryanodine receptors and the release of Ca^2+^ from endoplasmic reticulum stores, has been proposed as a potential underlying mechanism [14, 18–20].

In hippocampal neurons, ryanodine receptors (RyR) are expressed on the endoplasmic reticulum throughout the cell, including axons, dendrites and dendritic spines [21]. In situ hybridisation studies have revealed that type 3 ryanodine receptors (RyR3) are highly expressed, being indeed the predominant RyR subtype, in CA1 neurons of the rodent hippocampal formation, with a relatively lower level of expression in CA3 neurons [22–24]. Both CA1 and CA3 pyramidal neurons express also type 1 (RyR1) and type 2 (RyR2) receptors [22–24].

Ryanodine-sensitive calcium stores in CA1 pyramidal neurons contain a releasable pool of calcium that is maintained by calcium entry through voltage-gated calcium channels [25, 26]. Ca^2+^ influx evoked by either a single or multiple action potentials triggers RyR-mediated CICR from these stores, thereby increasing the overall magnitude of action potential-induced Ca^2+^ signals [27].

This action potential-induced Ca^2+^ elevation is essential to elicit the sAHP/sIAHP, but the actual contribution of RyR-mediated CICR to the generation of this afterpotential and K^+^ current in CA1 neurons is controversial in the existing literature. Some studies show that sAHP/sI_AHP_ is at least partly dependent on RyR-mediated CICR [14, 28–30], and particularly on RyR3 [31], while other studies confute any role of RyR-mediated CICR in the generation of sAHP/sI_AHP_ [17].

Here we have addressed the impact of RyR-mediated CICR on the amplitude of the sI_AHP_ at steady state and on its activity-dependent potentiation in rat and mouse hippocampal pyramidal neurons, and we have focused in particular on the role played by RyR3 in the regulation of sI_AHP_ in CA1 neurons by studying mice specifically lacking type 3 ryanodine receptors.%

## Materials and methods

### Ryanodine receptor type 3 deficient mice

Mice deficient in the ryanodine receptor type 3 gene (RyR3 -/-) were generated by homologous recombination, replacing exon 2 with a neomycin cassette, as described previously [32]. No Ryr3 protein was detected by Western blot analysis of brain tissue from RyrR -/- mice [32]. Mice were kept on a C57BL/6Jx129S4 background, and genotypes were confirmed by PCR on genomic DNA. All animal procedures were in accordance with the UK Animals (Scientific Procedures) Act 1986 and reviewed and approved by the UCL Animal Welfare and Ethical Review Body. Wild-type littermates (RyR3 +/+) from heterozygous crossings were used as internal controls. Experimenters remained blind to the genotype of RyR3 mice during experimentation and data analysis.

### Slice preparation

Acute slices were obtained from 21-28 days old Sprague Dawley rats or 3-5 months old RyR3 (RyR3 *-/-*; RyR3 *+/+*) mice. Animals were anaesthetized with isoflurane, decapitated and horizontal or transversal hippocampal slices (350 µm thick) were obtained using a vibratome (LeicaVT1000s, Leica, Germany) and were subsequently incubated in a humidified interface chamber at room temperature for ≥1 hr.

### Electrophysiology

Tight-seal whole-cell patch clamp recordings were obtained from CA1 pyramidal neurons using the “blind patching technique” [33]. Experiments were conducted either with an EPC9 or an EPC10 amplifier (HEKA, Germany) controlled by Pulse or PatchMaster software for data acquisition (HEKA, Germany). Slices were perfused in a submerged recording chamber with a constant flow of 2-2.5 ml/min with carbogen-bubbled ACSF containing (in mM: 125 NaCl, 1.25 KCl, 2.5 CaCl_2_, 1.5 MgCl_2_, 1.25 KH_2_PO_4_, 25 NaHCO_3_, and 16 D-glucose) and the sI_AHP_ was & recorded at room temperature (22±1°C) with patch pipettes made of borosilicate glass (Hilgenberg, Germany), with a resistance of 4.5-7.5 MOhm when filled with intracellular solution. For recordings from rat CA1 pyramidal neurons, the pipette solution used contained (in mM): 135 K-gluconate, 10 KCl, 10 HEPES, 2 Na_2_ATP, 0.4 Na_3_GTP 0.4 and 1 MgCl_2_. For recordings from mouse CA1 pyramidal neurons, the pipette solution contained (in mM): 135 K-MeSO_4_, 10 KCl, 10 HEPES, 2 Na_2_ATP, 0.4 Na_3_GTP and 1 MgCl_2_. The pH was adjusted to 7.2-7.3 with KOH and the osmolarity of the intracellular solution was between 280-290 mOsm/kg. Only cells with a resting membrane potential !-55 mV and a series resistance !30 MOhm, not changing by more than 25% in the course of the experiment, were included in this study. Voltage values reported were not corrected for the liquid junction potential that was -11 mV (intracellular K-gluconate solution) and -5 mV (intracellular K-MeSO_4_ solution).

Action potentials were elicited by 40-100 pA, 1 s-long current injections repeated and increased by further 40-100 pA every 20 s. Data were filtered at 3 kHz and sampled at 12.5 kHz. Series resistance, input resistance and membrane time constant (τ) of CA1 pyramidal neurons were measured in response to 100 ms-long voltage steps of -5mV from a holding potential of -50 mV; data were filtered at 5 kHz and sampled at 20 kHz.

The sI_AHP_ was measured as an outward current and elicited by stepping to +10 mV for 100 ms from a holding potential of -50 mV every 30 seconds to activate voltage-gated Ca^2+^ channels and obtain Ca^2+^ influx necessary to activate the sI_AHP_. Traces data were sampled at 1 kHz and filtered at 250 Hz. In all voltage-clamp recordings we added to the superfusing ACSF tetrodotoxin (TTX, 0.5 µM) to block voltage-gated sodium channels, and tetraethylammonium (TEA, 1 mM) to block a subset of voltage-gated potassium channels and increase calcium influx and thereby the calcium dependent sI_AHP_. In recordings from mouse CA1 pyramidal neurons, also d-tubocurarine hydrochloride (dTC, 100 µM) was added to the ACSF to inhibit SK channels. 10 µM Ryanodine was applied either for ∼15 minutes after sI_AHP_ had reached a steady state amplitude (i.e. upon completion of the potentiation phase) or for the whole duration of the whole-cell recording to study its impact on the sI_AHP_ potentiation. In some experiments (Rp)-Adenosine-3’,5’- monophosphorothioate (Rp-cAMPS, 500 µM) was added to the intracellular solution to inhibit protein kinase A (PKA).

### Data analysis and statistics

All experiments were analysed using the software Pulsefit (HEKA, Germany) and IGOR Pro (Wavemetrics, USA) with the support of Neuromatic [34]. The amplitude of the sI_AHP_ was measured 700-1000 ms after the end of the command pulse, when possible contamination by other, faster outward currents are negligible [35]. The charge transfer was determined as the area under the curve starting from the sI_AHP_ peak until full decay had occurred. The deactivation time constant (τ_decay_) was obtained by fitting a mono-exponential function to the data points.

The amplitudes of 2-3 traces around a given time-point or before and after drug application were averaged to quantify potentiation or drug effects, as shown in summary bar diagrams and box and whisker plots.

The activity-dependent potentiation (run-up) was calculated by normalising the sI_AHP_ amplitude to the amplitude recorded at 0 min. The corresponding time constant of sI_AHP_ potentiation (τ_potentiation_) was calculated by fitting a mono-exponential function to the first 15 minutes of the run-up phase. To quantify the potentitation, sI_AHP_ amplitudes were measured at the start (0 min) and the end (15 min) of the potentiation, and used to calculate the ratio (sI_AHP_ (15 min)/sI_AHP_ (0 min)).

All graphs were created using Prism 6 (GraphPad, USA), which was also used for the statistical analysis, together with InStat (GraphPad, USA). In box-and-whiskers plots, the boxes extend from the 25th to 75th percentiles, the lines in the middle of the boxes represent the median, the circles correspond to the mean and the whiskers stretch to the smallest and largest values in each data set. Mean and standard error of the mean (mean ± SEM) were used to describe the results of statistical analysis, as shown in bar diagrams and for the data points in the graphs showing the (relative (%) time-courses of potentiation. When not otherwise specified, for data with normal distributions, paired or unpaired t-tests or ANOVA were used for comparisons; when data were not normally distributed, appropriate non-parametric tests or corrections were employed to calculate statistical significance, as specified in the text.

### Drugs and solutions

Ryanodine was obtained from Calbiochem (Millipore, Heartfordshire, UK) and Alomone Labs (Israel); cyclopiazonic acid (CPA) from Alomone Labs (Israel); caffeine from Calbiochem (Millipore, Heartfordshire, UK); KMeSO_4_ from Fisher Scientific (Loughborough, UK); tetrodotoxin (TTX) citrate-free from Latoxan (Rosans, France); d-tubocurarine hydrochloride (dTC), Na_2_ATP, Na_3_GTP and tetraethylammonium (TEA) from Sigma-Aldrich (Dorset, UK); (Rp)-Adenosine-3’,5’- monophosphorothioate (Rp-cAMPS) (BioLog Life Science Institute, Germany). All other salts and chemicals were obtained from Sigma-Aldrich or VWR International Ltd. Drugs were dissolved in water or DMSO, stored at +4°C or -20°C, and bath applied in the perfusing ACSF.)

## Results

### RyR-mediated CICR affects sI_AHP_ and its potentiation in rat CA1 pyramidal neurons

The impact of Ca^2+^-induced Ca^2+^ release (CICR) on the sI_AHP_ in rat CA1 pyramidal neurons was studied with a set of pharmacological tools targeting ryanodine receptor (RyR)-mediated CICR. The effects of these compounds on both the steady-state amplitude and the activity-dependent potentiation of the sI_AHP_ were assessed.

Whole cell voltage-clamp recordings were obtained from 72 CA1 pyramidal neurons in rat hippocampal slices. sI_AHP_ was elicited as a tail current following an inward Ca current activated by a 100 ms long depolarizing voltage step from -50 mV to +10 mV which was repeated every 30 s. The mean resting membrane potential in all measured cell was -59.5 ± 0.4 mV (n = 72) and the mean amplitude of sI_AHP_ at steady-state was 59.3 ± 3.7 pA (n = 72)

### Caffeine enhances sI_AHP_ in rat CA1 pyramidal neurons

The methylxanthine caffeine binds to the ryanodine receptors and, when used at a low concentration (0.5 mM) [36], enhances the increase in intracellular Ca^2+^ due to Ca^2+^ release from the ryanodine-sensitive stores [26, 37, 38]. Therefore, we tested whether the application of caffeine at this concentration affected the sI_AHP_.

A biphasic effect of caffeine (0.5 mM) on the sI_AHP_ amplitude (Fig 1A-E) was observed in 16 cells. In these cells, the sI_AHP_ amplitude first increased by ∼25% (P < 0.0001), followed by a decrease by ∼25% when compared to the amplitude of the current preceding caffeine application. In a further six cells the same increase in the sI_AHP_ amplitude was observed, but the recording condition became unstable before the time point to estimate the amplitude reduction was reached.

**Fig 1.**
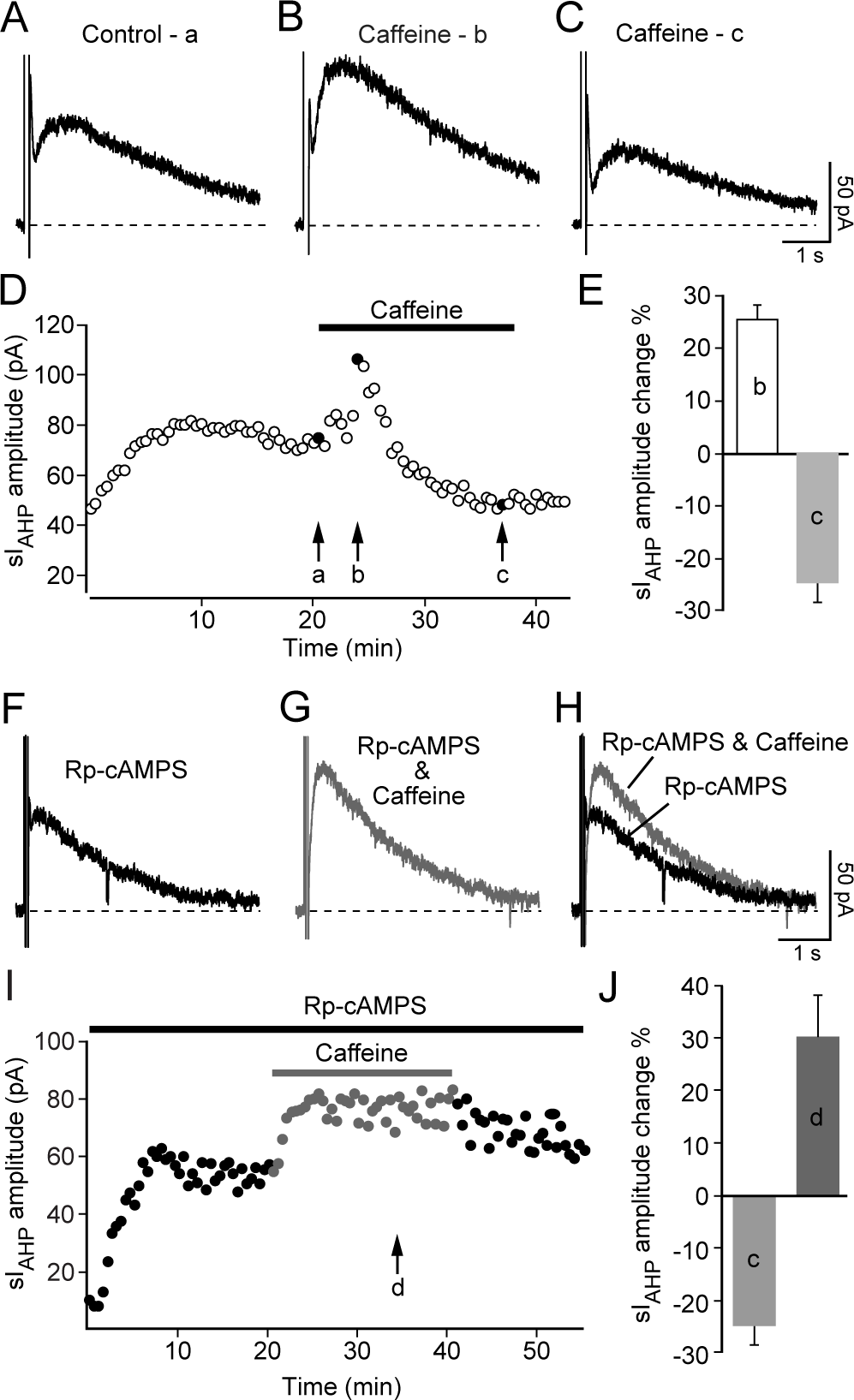
Effect of caffeine on sI_AHP_ in rat CA1 neurons. **(A)** Representative sI_AHP_ trace at steady state. **(B)** sI_AHP_ trace maximally enhanced after 2.5 min into the application of caffeine (0.5 mM). **(C)** sI_AHP_ reduction after continuous application of caffeine. **(D)** Time course of action of caffeine (0.5 mM) on the sI_AHP_ amplitude. Traces **(A-C)** labelled as **a**, **b** and **c** were taken from the same rat CA1 pyramidal neuron. **(E)** Summary bar diagram, showing that the sI_AHP_ amplitude first increased by 25.3 ± 2.9% (**b**: n = 22; P < 0.0001), followed by a decrease by 24.5 ± 3.6% (**c**: n = 16; P < 0.0001) when compared to the current amplitude preceding caffeine application. **(F)** Representative sI_AHP_ trace recorded at steady state but with the PKA inhibitor Rp-cAMPS (500 µM) applied intracellularly. **(G)** sI_AHP_ current increase in the presence of Rp-cAMPS and after ∼17.5 min application of caffeine (0.5 mM), a time point comparable to (**c**) in **(A)**. **(H)** Overlay of the sI_AHP_ traces in **F** and **G**. (**I)** Time course of action of caffeine (0.5 mM) in the presence of Rp-cAMPS; (**d**) indicates the trace shown in **G**. **(J)** Bar diagram comparing the relative (%) changes in sI_AHP_ amplitude in response to caffeine application in the absence (**c**; of panel **E**) and presence of the PKA inhibitor Rp-cAMPS (**d**). sI_AHP_ amplitude increased by 29.8 ± 7.9% (**d**: n = 4; P = 0.03) in the presence of Rp-cAMPS after caffeine application when compared to the sI_AHP_ current amplitude preceding caffeine application. The difference between the effect of caffeine with or without Rp-cAMPS is highly significant (P < 10^-6^).

Beside enhancing CICR from intracellular Ca^2+^ stores, caffeine is also an effective inhibitor of phosphodiesterases [39, 40]. The inhibition of phosphodiesterases leads to an increase of intracellular cAMP known to inhibit the sI_AHP_ through activation of protein kinase A (PKA) [41, 42].

Therefore, if the observed decrease of the sI_AHP_ amplitude during prolonged caffeine application was due to the elevation of intracellular cAMP and consequent activation of PKA, then this effect should be prevented by inhibition of PKA. This hypothesis was tested by applying the specific PKA-inhibitor Rp-cAMPS (500 #M), which competitively inhibits cAMP-binding sites on the regulatory subunits of PKA [43], through the patch pipette and allowing its diffusion into the cell before the application of caffeine. In the presence of Rp-cAMPS, caffeine increased sI_AHP_ by a comparable percentage (29.8 ± 7.9%) as in the absence of Rp-cAMPS (Fig 1F-J). However, this increase of sI_AHP_ amplitude was not followed by a decrease, supporting the hypothesis that indeed the decrease in sI_AHP_ was due to activation of PKA.

These results show that enhancement of intracellular Ca^2+^ levels caused by stimulation of ryanodine-sensitive stores by caffeine leads to an increase in sI_AHP_ amplitude. While this observation establishes a link between Ca^2+^ released from intracellular stores and sI_AHP_, it does not address the question as to whether CICR contributes to the activation of sI_AHP_ when the current is elicited in response to depolarizing stimuli that increases intracellular Ca^2+^ by activating voltage-gated Ca^2+^ channels.

### Ryanodine reduces the sI_AHP_ amplitude at steady-state in rat CA1 pyramidal neurons

To address the question of the concurrent involvement of RyR-mediated CICR in the generation of the sI_AHP_ elicited by activation of voltage-gated Ca channels in response to depolarizing stimuli ryanodine was used. At a concentration of 10 µM ryanodine binds to the RyR receptor and keeps it in an open subconducting state thereby depleting the ryanodine sensitive Ca^2+^ store [26, 44].

Ryanodine (10 µM) was bath-applied once the sI_AHP_ had reached a stable amplitude, in order to assess its effect on the current at steady-state (Fig 2A-D). In response to ryanodine, the amplitude of the sI_AHP_ decreased by ∼23%, from 70.0 ± 2.8 pA to 53.5 ± 2.5 pA (n = 5; P = 0.004; Fig 2B-E).

**Fig 2.**
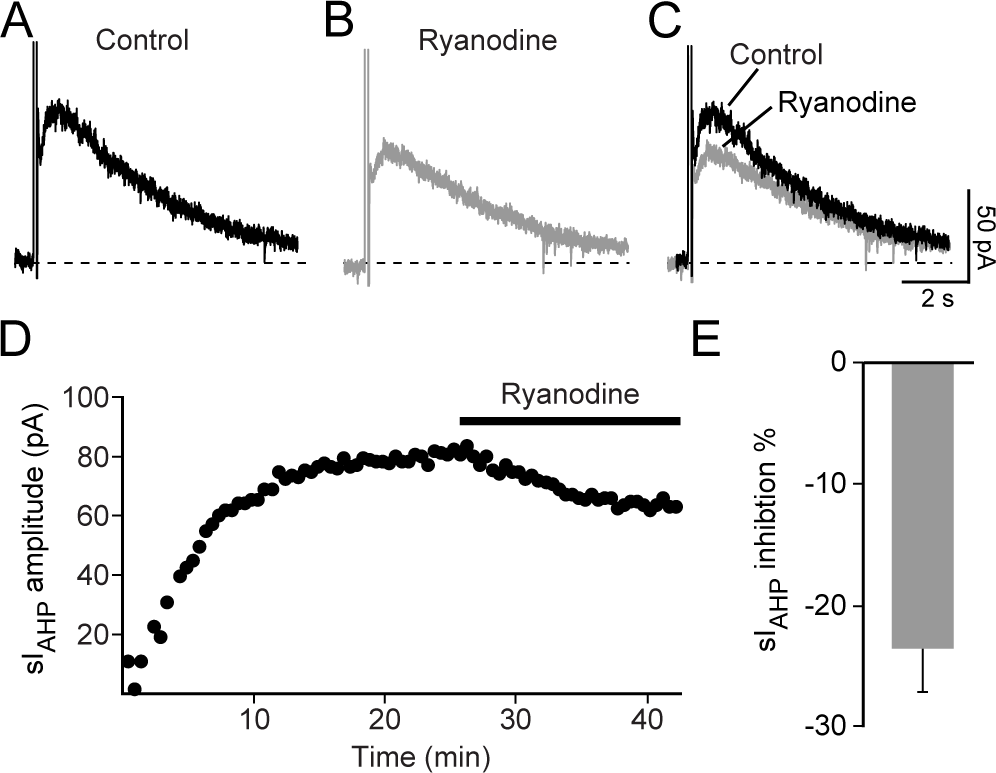
Effect of ryanodine on the sI_AHP_ measured at steady state in rat CA1 pyramidal neurons. **(A)** Representative sI_AHP_ trace measured at steady state. **(B)** Reduction of the sI_AHP_ upon application of ryanodine (10 µM). (**C)** Superimposed traces before and after application of ryanodine. (**D)** Time course of action of ryanodine (10 µM) on the sI_AHP_ amplitude in the same cell as for the traces shown in panels **A-C**. **(E)** Summary bar diagram showing that ryanodine decreases the amplitude of the sI_AHP_ by 23.3 ± 3.7% (n = 5; P = 0.03).

This result suggests that in rat CA1 pyramidal neurons CICR from ryanodine-sensitive stores makes a small, but significant contribution to the generation of the sI_AHP_ primarily evoked in response to Ca^2+^ influx through voltage-gated Ca^2+^ channels activated by depolarizing stimuli.

### Inhibition of CICR by ryanodine hinders the activity-dependent potentiation of sI_AHP_ in rat CA1 pyramidal neurons

Repeated activation of CA1 pyramidal neurons by depolarizing pulses induces a reduction in excitability, most likely as a consequence of an increased sI_AHP_ [13, 15]. The activity-dependent potentiation or “run-up” [14, 17], a phase resulting in a substantial and sustained increase of sI_AHP_, has been attributed to an increase in intracellular Ca^2+^ [14, 16]. Therefore we addressed the question as to whether ryanodine-dependent CICR contributes to the sI_AHP_ potentiation in whole-cell patch clamp recordings.

A run-up phase of the sI_AHP_, lasting in average 15 min, was consistently observed in our recordings (e. g. Fig 1D, 1I and Fig 2D). To analyse the run-up phase in more detail, TTX (0.5 µM) and TEA (1 mM) were added to the bath solution at least 3 minutes before recording the first sI_AHP_ trace, in order to exclude potential contributions induced by the sodium and potassium channel blockers to the sI_AHP_ amplitude increase. In control experiments (0.2 % DMSO), the sI_AHP_ amplitude increased from 23.1 ± 5.1 pA to 79.0 ± 7.7 pA (Fig 3A-C and J-K), a nearly 4 fold increase after 15 minutes (n = 5; Fig L, paired t-test: P = 0.0017), with 75% of the potentiation occurring already within the first 3 minutes (Fig 3 J and L).

**Fig 3.**
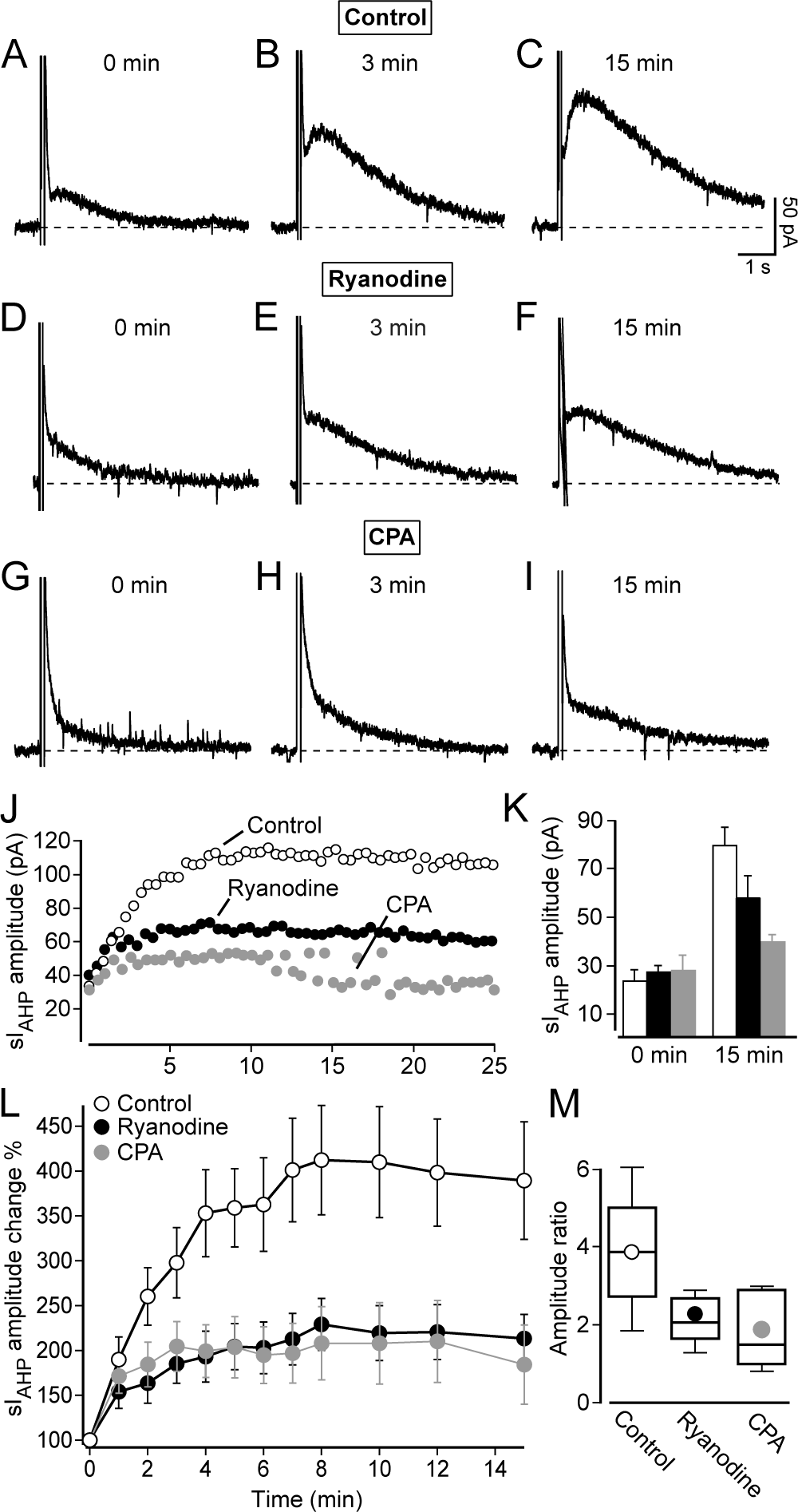
CICR contributes to the activity-dependent potentiation of sI_AHP_ in rat CA1 pyramidal neurons. (**A-I)** Representative current traces at 0 min, 3 min and 15 min show the potentiation of the sI_AHP_. (**A-C)** Control traces recorded in the presence of 0.2% DMSO. (**D-F)** Reduced potentiation of the sI_AHP_ in the presence of 10 #M ryanodine. (**G-I)** Reduced potentiation of the sI_AHP_ in the presence of 50 #M CPA. (**J)** Superimposed time-courses of three representative cells comparing the different degrees of the potentiation of sI_AHP_ amplitude under control conditions, 10 µM ryanodine and 50 µM CPA. (**K)** Bar chart summarizing the sI_AHP_ amplitude at the beginning of the recording (0 min) and the end of the recording (15 min), (control, 0 min: 23.1 ± 5.1 pA; 15 min: 79.0 ± 7.7 pA; n = 5; paired t-test: P = 0.0017; 10 µM ryanodine, 0 min: 27.4 ± 2.6 pA; 15 min: 58.2 ± 9.3 pA; n = 5; paired t-test: P = 0.02; 50 µM CPA; 0 min: 27.5 ± 6.6 pA; 15 min: 39.2 ± 3.1 pA; n = 5; paired t-test: P = 0.12). (**L)** Summary of the relative (%) time-courses of potentiation of sI_AHP_ measured in all cells (n = 5 for each condition) under control conditions (0.2% DMSO), in 10 µM ryanodine and in 50 µM CPA. Reduction of the sI_AHP_ amplitude potentiation in the presence of ryanodine (two-way ANOVA with post-hoc Bonferroni test: P < 0.0001) and of CPA (two-way ANOVA with post-hoc Bonferroni test: P < 0.0001). (**M)** Ratio of the sI_AHP_ amplitude measured at 0 and 15 min under control conditions (mean ± SEM: 3.9 ± 0.7; n = 5), in ryanodine (2.2 ± 0.3; n = 5; P = 0.04) and in CPA (1.9 ± 0.4; n = 5; P = 0.03).

Next we tested whether the run-up phase of sI_AHP_ depends on CICR from ryanodine-sensitive stores. Therefore 10 µM ryanodine was applied from the very beginning of the recording. Also under these conditions the sI_AHP_ amplitude increased significantly within the first 15 min of the recording from 27.4 ± 2.6 pA to 58.2 ± 9.3 pA (Fig 3 D-F; J and L, n = 5, paired t-test: P = 0.02).

Although the sI_AHP_ amplitude increased within 15 min under control conditions as well as in the presence of ryanodine, the extent of the sI_AHP_ amplitude potentiation was significantly reduced in the presence of ryanodine (Fig 3L; n = 5; two-way ANOVA with post-hoc Bonferroni test: P < 0.0001).

The ratio of the sI_AHP_ amplitude measured at 0 and 15 min under control conditions and in the presence of ryanodine shows that the sI_AHP_ potentiation has been reduced by roughly 50% by ryanodine (Fig 3M). The extensive attenuation in the sI_AHP_ run-up observed in the presence of ryanodine suggests that Ca^2+^ released by RyR-mediated CICR contributes substantially in mediating the activity-dependent potentiation of sI_AHP_.

### Inhibition of Ca^2+^ store refilling reduces the activity-dependent potentiation of sI_AHP_ in rat CA1 pyramidal neurons

The sI_AHP_ activity-dependent potentiation depends on RyR-mediated CICR, as suggested by its reduction by ryanodine. If this is the case, the run-up should be affected when the Ca^2+^-reuptake into the endoplasmic reticulum compartment is inhibited. Cyclopiazonic acid (CPA) is an endoplasmic reticulum Ca^2+^-ATPase blocker that prevents the refilling of depleted Ca^2+^ stores in CA1 pyramidal neurons [26]. Consequently, in the presence of CPA no further Ca^2+^ can be released from intracellular stores after an initial release.

CPA was used in the next set of experiments to see whether the suppression of the Ca^2+^ store refilling resulted in a concomitant reduction of the activity-dependent potentiation. CPA (50 µM) was added together with TTX (0.5 µM) and TEA (1 mM) to the bath solution at least 3 minutes before recording the first sI_AHP_ trace. Although the current increased slightly (from 27.5 ± 6.6 pA at 0 min to 39.2 ± 3.1 pA at 15 min; n = 5; Fig 3 G-I and J-K), no significant increase in the sI_AHP_ amplitude was observed within the first 15 min (Fig 3 K and L) of the recording in the presence of CPA (paired t-test: P = 0.12; n = 5) showing that preventing the refilling of the Ca^2+^ stores by CPA reduces the activity-dependent potentiation as efficiently as depleting the stores by 10 µM ryanodine. Overall the extent of the sI_AHP_ amplitude potentiation was significantly reduced in the presence of CPA (two-way ANOVA with post-hoc Bonferroni test: P < 0.0001; n = 5; Fig 3L). The conspicuous reduction by around 50 % (Fig 3M) of the sI_AHP_ amplitude ratio in the presence of CPA further illustrates the impact of ryanodine-sensititve store depletion on the sI_AHP_ potentiation.

Taken together, the results obtained with ryanodine and CPA demonstrate that RyR-mediated CICR is an essential mechanism underlying the activity-dependent potentiation of sI_AHP_ in rat CA1 pyramidal neurons.

### Features of sI_AHP_ in CA1 pyramidal neurons from mice lacking type 3 ryanodine receptor (RyR3) and wildtype littermates

The results shown so far (Fig 1-3) imply that RyR-mediated CICR contributes to the maintenance and potentiation of sI_AHP_ in rat CA1 pyramidal neurons. This leads to the question as to which of the three existing receptor subtypes (RyR1, RyR2 and RyR3) is involved in this process, since ryanodine binds to all three. Both RyR2 and RyR3 are expressed in the CA1 layer, with RyR3 being the most abundant subtype in CA1 pyramidal cells, followed by RyR2 [23]. Moreover, RyR3 was proposed to be specifically responsible for triggering sI_AHP_, as intracellular application of anti-RyR3 antibodies strongly reduced the current amplitude in mouse CA1 pyramidal neurons [31].

Therefore, we next investigated whether RyR3 plays a role in the generation and maintenance of sI_AHP_ and is responsible for the activity-dependent potentiation of sI_AHP_ by using knock-out mice lacking RyR3 [32].

Before studying how the absence of the RyR3 receptor affects the generation, maintenance and activity-dependent potentiation of the sI_AHP_, we investigated whether the loss of RyR3 changed the passive membrane and firing properties of mouse CA1 pyramidal neurons. The mean resting membrane potential of CA1 pyramidal neurons was not affected by the lack of RyR3 (RyR3 $/$) = - 66.8 ± 0.6 mV; n = 61; (RyR3 +/+) = -67.0 ± 0.6 mV; n = 60) (P = 0.9, Mann-Whitney test). Also the membrane time constant of CA1 pyramidal neurons in wild-type littermates (RyR3 +/+, 7.5 ± 0.3 ms, n = 60) did not change compared to RyR3 $/$ mice (7.5 ± 0.3 ms, n = 61; P = 0.7, Mann-Whitney test). However, the input resistance CA1 pyramidal neurons was significantly lower in RyR3 $/$ (204.1 ± 7.5 M%, n = 61) compared to RyR3 +/+ littermates (251.2 ± 8.6 M%, n = 60) (P < 0.0001). Conversely, the membrane capacitance of CA1 pyramidal neurons from RyR3 $/$ mice (41.6 ± 2.9 pF, n = 61) was higher compared to RyR3 +/+ littermates (32.9 ± 2.1 pF, n = 60, P = 0.03, Mann-Whitney test).

When comparing the firing rates of CA1 pyramidal neurons from RyR3 +/+ and RyR3 $/$ littermates in response to increasing current injections, CA1 neurons recorded from RyR3 +/+ mice fired more action potentials, at a higher frequency compared to RyR3 $/$ CA1 neurons (P = 0.004, two-way ANOVA with Bonferroni’s test) (Fig 4A-C).

**Fig 4.**
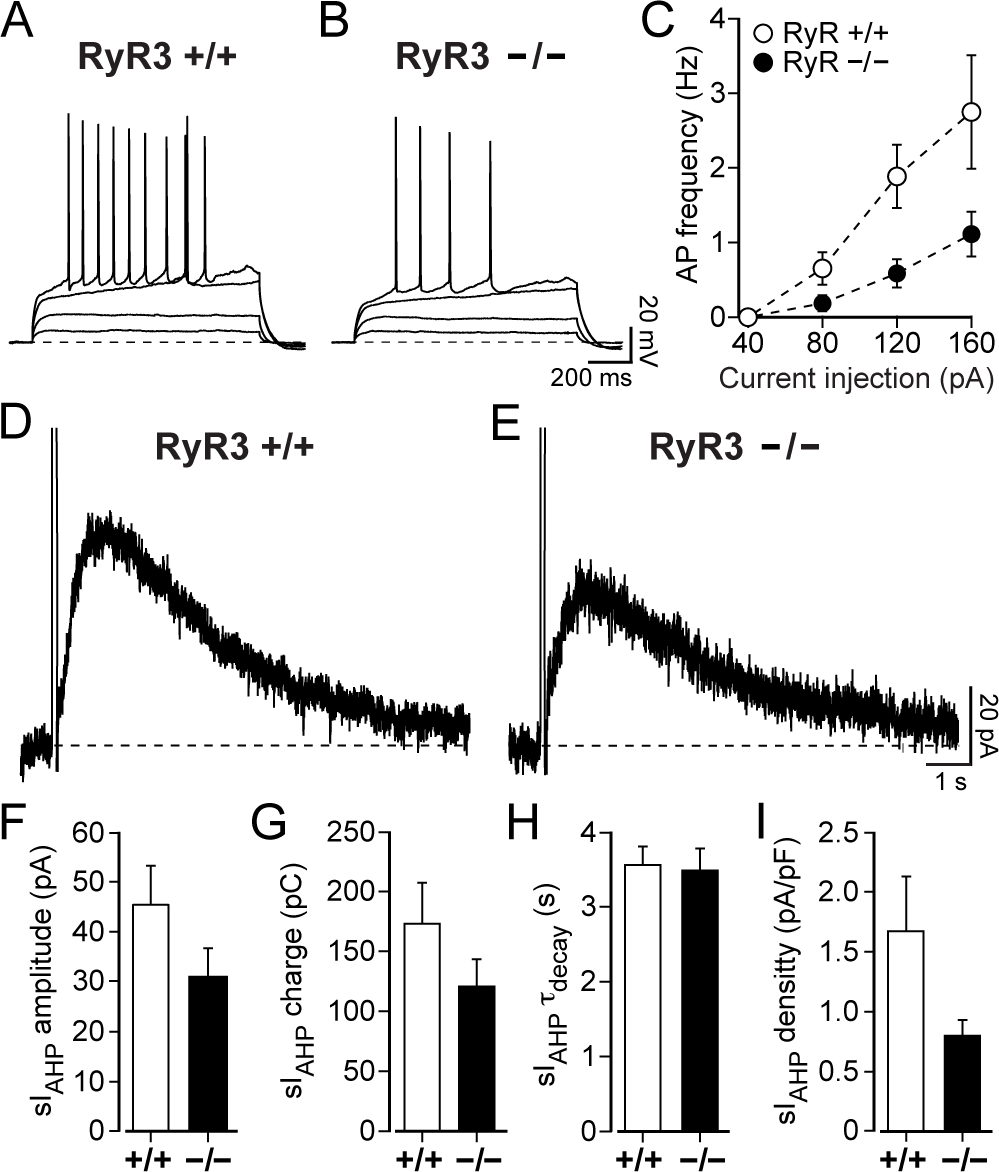
Lack of type 3 ryanodine receptor (RyR3) affects the firing properties and sI_AHP_ current density in mouse CA1 pyramidal neurons. (**A-B)** action potentials were induced by current injections from 40 pA to 160 pA (in 40 pA step-increments) for 1 s in RyR3 +/+ and RyR3 -/- CA1 pyramidal neurons. The membrane resting potential was -68 mV (A) and -69 mV (B). (**C)** Action potential frequency was higher in RyR3 +/+ compared to RyR3 -/- CA1 pyramidal neurons at the same stimulation strength (P = 0.004, two-way ANOVA with Bonferroni’s test). At 40 pA action potentials could not be elicited in either RyR3 +/+ or RyR3 -/- CA1 pyramidal neurons. The action potential frequency did not differ significantly (P = 0.5) for current injections at 80 pA for RyR3 +/+ (0.7 ± 0.2 Hz, n = 32) and RyR3 -/- (0.2 ± 0.1 Hz, n = 37). However, at higher current injections action potential frequencies were substantially different (120 pA: RyR3 +/+ = 1.9 ± 0.4 Hz, n = 27 vs RyR3 -/- = 1.1 ± 0.2 Hz, n = 34, P = 0.0003; and 160 pA RyR3 +/+ = 2.8 ± 0.8 Hz, n = 12 vs RyR3 -/- = 1.1 ± 0.3 Hz, n = 26, P = 0.001). (**D-E**) Representative sI_AHP_ traces obtained from RyR3 +/+ and RyR3 -/- mouse CA1 pyramidal neurons. (**F-I**) Properties of the sI_AHP_ recorded at steady-state in RyR3 +/+ and RyR3 -/- mice ∼15 minutes after the onset of the recording. (**F**) No significant difference was observed for current amplitude (RyR3 +/+ = 42.5 ± 8.1 pA, n = 10 vs RyR3 -/- = 30.5 ± 6.1 pA, n = 7; P = 0.2). (**G)** Similarly, there was no significant difference for sI_AHP_ charge transfer (RyR3 +/+ = 172.9 ± 34.6 pC, n = 10 vs RyR3 -/- = 120.6 ± 22.8 pC, n = 7; P = 0.3) and (**H)** the deactivation time constant of sI_AHP_(RyR3 +/+ = 3.6 ± 0.3 s, n = 10 vs RyR3 -/- = 3.5 ± 0.3 s, n = 7; P = 0.9). (**I)** sI_AHP_ current density was reduced in RyR3 -/- neurons (RyR3 -/- = 0.8 ± 0.1 pA/pF, n = 7 vs RyR3 +/+ = 1.7 ± 0.5 pA/pF, n = 10; P = 0.03, Mann-Whitney test).

In whole cell voltage-clamp recordings from CA1 pyramidal neurons in RyR3 +/+ and RyR3 $/$ littermates, the sI_AHP_ was elicited with the same pulse protocol previously used to stimulate rat CA1 pyramidal neurons (100 ms long depolarizing voltage steps from -50 mV to +10 mV, repeated every 30 s). The sI_AHP_ was present in both RyR3 +/+ and RyR3 $/$ (Fig 4D and E) CA1 pyramidal neuron at the end of the run-up phase, 15 min from the onset of the recording. At this time point the sI_AHP_ amplitude (Fig 4F; P = 0.2), charge transfer (Fig 4G; P = 0.3), and deactivation time constant (Fig 4H; P = 0.9) were similar in CA1 pyramidal neurons from RyR3 +/+ and RyR3 $/$ mice. The sI_AHP_ density (Fig 4I), calculated by normalising the sI_AHP_ amplitude to the membrane capacitance, was reduced in CA1 pyramidal neurons of RyR3 $/$ compared to neurons in RyR3 +/+ mice (P = 0.03, Mann-Whitney test), as expected due to the increased membrane capacitance of RyR3 $/$ neurons.

These results show that the absence of RyR3 did not prevent the generation and maintenance of the sI_AHP_ in CA1 pyramidal neurons, and fundamental sI_AHP_ properties were not different in RyR3 $/$ and RyR3 +/+ (Fig 4D-H), with the exception of sI_AHP_ density, which was lower in RyR3 $/$ than in RyR3 +/+ CA1 pyramidal neurons (Fig 4I).

### Ryanodine reduces the sI_AHP_ amplitude at steady-state in RyR3 +/+ and RyR3 !/! CA1 pyramidal neurons

The next step was to study the effect of CICR inhibition by ryanodine on the sI_AHP_ measured at steady-state in wild type mice and mice lacking RyR3. At the end of the run-up phase, once the sI_AHP_ was fully potentiated, normally ∼15 minutes after the beginning of the recording (Fig 5G), application of ryanodine (10 #M) to RyR3 +/+ CA1 pyramidal cells reduced the amplitude of the sI_AHP_ by 39.0 ± 6.9% (n = 7, Fig 5A-C, G and I). The sI_AHP_ reduced in amplitude over a period of 15- 20 minutes (Fig 5G), comparable with our observation in rat CA1 pyramidal neurons (Fig 2D). Thus, ryanodine application has similar effects on the sI_AHP_ measured at steady state in mouse and rat CA1 pyramidal neurons.

**Fig 5.**
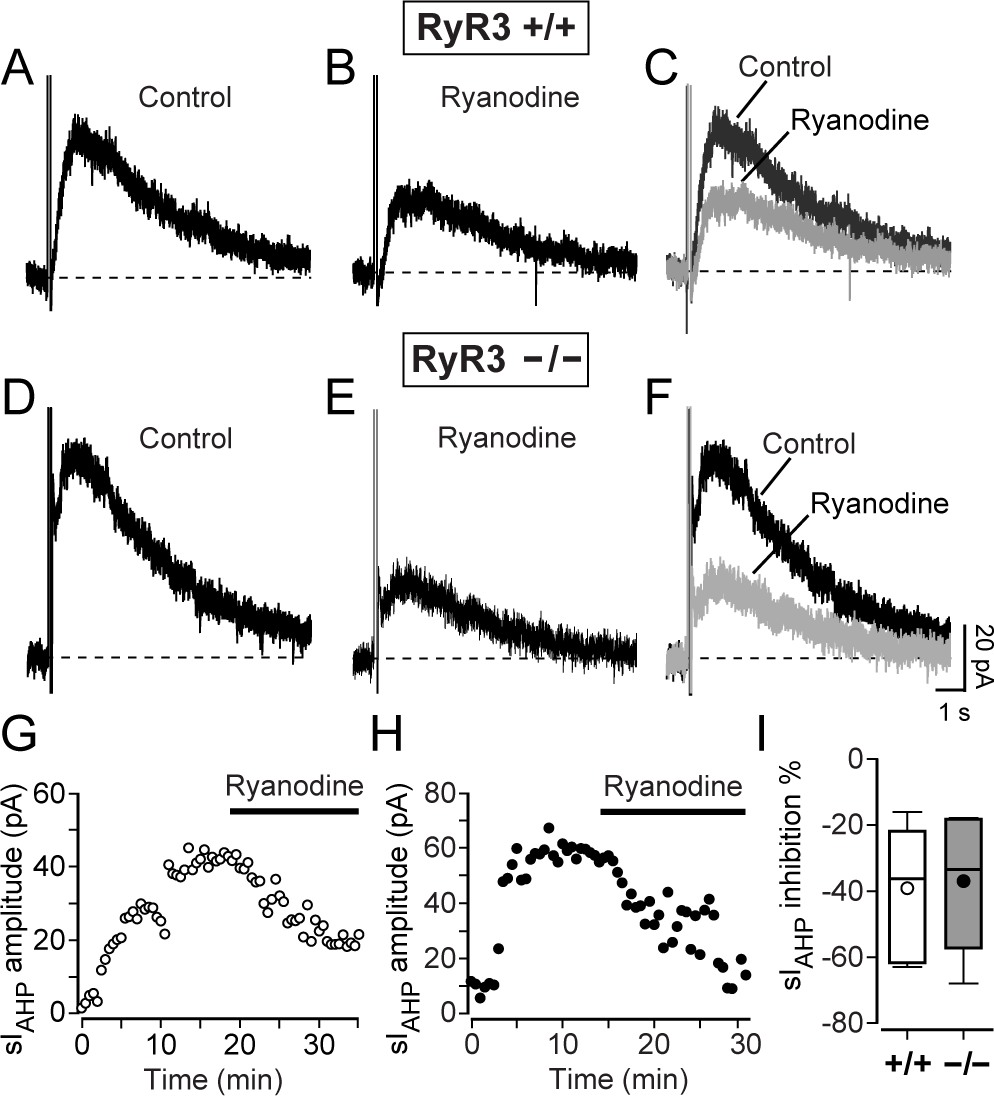
Inhibition of CICR by ryanodine reduced the sI_AHP_ amplitude at steady state in RyR3 +/+ and RyR3 -/- mouse CA1 pyramidal neurons. Representative sI_AHP_ traces obtained from RyR3 +/+ (**A**) and RyR3 -/- (**D**) mouse CA1 pyramidal neurons at steady-state, at the end of the run-up phase. Current traces from the same neurons 15 minutes after the application of 10 µM ryanodine (RyR3 +/+ (**B**) and RyR3 -/- (**E**)). (**C** and **F)** Superimposed traces showing the reduction of the sI_AHP_ amplitude. (**G**) Time course of sI_AHP_ amplitude in the same RyR3 +/+ neuron as shown in A-C. (**H**) Time course of the sI_AHP_ amplitude in the same RyR3 -/- neuron as shown in D-F. (**I)** Overall decrease in the sI_AHP_ amplitude summarized in a box-and-whiskers plot, showing comparable reduction by ryanodine in RyR3 +/+ (mean ± SEM: 39.0 ± 6.9 %; median = 36.2 %, n = 7) and RyR3 -/- neurons (mean ± SEM: 36.9 ± 9.4%; median = 33.4 %, n = 5) (P = 0.9).

The same experimental conditions were used to study the effect of ryanodine application on RyR3 $/$ CA1 pyramidal cells. After the sI_AHP_ was fully potentiated and reached a steady state amplitude (Fig 5D and H), 10 #M ryanodine was applied. Ryanodine application reduced the amplitude of the sI_AHP_ by 36.9 ± 9.4 % (n = 7, Fig 5E-F, H and I). The reduction in sI_AHP_ amplitude caused by ryanodine was comparable in RyR3 +/+ and RyR3 $/$ CA1 pyramidal neurons (P = 0.9) (Fig 5I). These results show that inhibition of CICR by ryanodine has a similar impact on sI_AHP_ measured at steady state in the presence and in the absence of type 3 ryanodine receptors.

### Lack of RyR3 affects the activity-dependent potentiation of sI_AHP_ in mouse CA1 pyramidal neurons

In rat CA1 neurons, sI_AHP_ undergoes an activity-dependent potentiation that is sensitive to RyR-mediated CICR (Fig 3) [14]. We therefore investigated whether RyR3 played a specific role in mediating the sI_AHP_ potentiation by measuring and comparing it in RyR3 +/+ and RyR3 $/$ CA1 pyramidal neurons. As in the recordings before, when characterizing the run-up phase in rat CA1 pyramidal neurons, the inhibitors for sodium and potassium channels were added to the bath solution at least 3 minutes before recording the first sI_AHP_ trace. The sI_AHP_ started as a small current and increased to a larger amplitude by 15 minutes in RyR3 +/+ (one-way ANOVA, P < 0.05) and RyR3 $/$ (one-way ANOVA, P < 0.0001) CA1 pyramidal cells (Fig 6A, B and D). The starting sI_AHP_ amplitudes in RyR3 +/+ and RyR3 $/$ CA1 neurons were not significantly different (P = 0.07, Mann-Whitney test) (Fig 6C), despite a tendency towards larger initial currents in RyR3 $/$ neurons. At the end of the run-up phase, at 15 minutes, the steady-state sI_AHP_ amplitude was also similar in CA1 neurons from RyR3 +/+ and RyR3 $/$ mice (P = 0.2) (Fig 6C).

**Fig 6.**
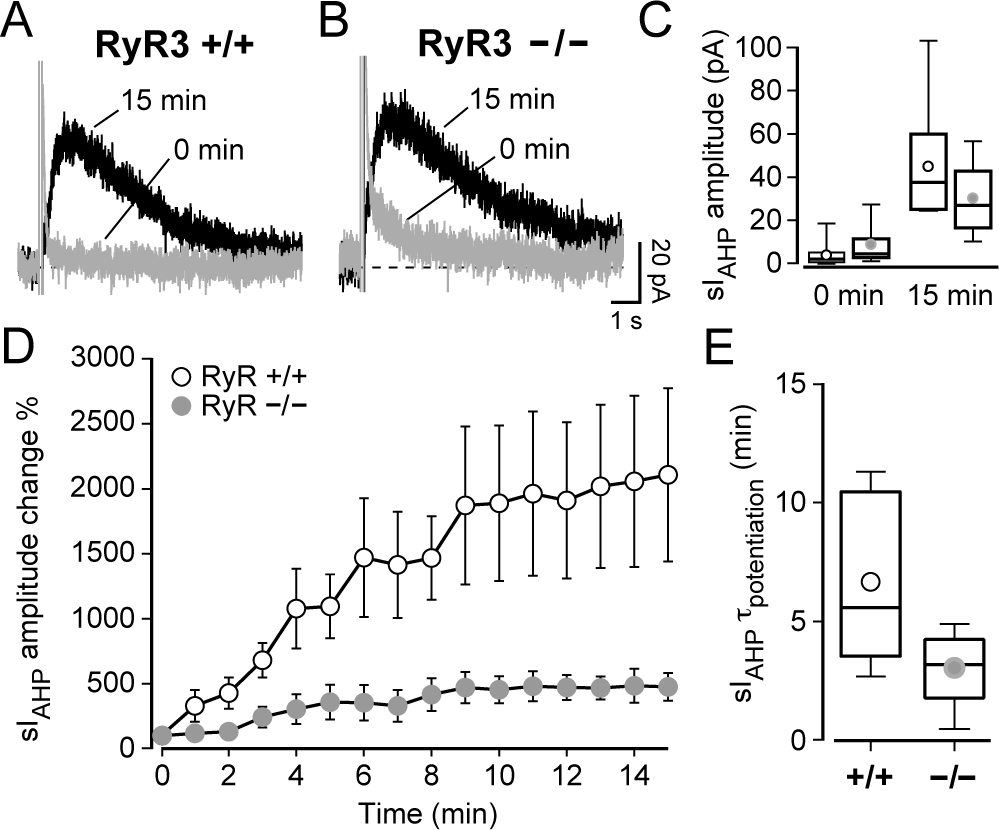
CA1 pyramidal neurons lacking type 3 ryanodine receptor (RyR3 -/-) have a faster and reduced activity-dependent potentiation of sI_AHP_. (**A, B)** Superimposed traces of the sI_AHP_ recorded at 0 and 15 minutes from RyR3 +/+ and RyR3 -/- CA1 pyramidal neurons. (**C)** Box-and-whiskers plot summarizing the sI_AHP_ amplitudes recorded before the run-up phase (0 min) and at the end of potentiation (15 min). At 0 min the sI_AHP_ peak amplitude in RyR3 +/+ (mean ± SEM: 4.2 ± 1.8 pA; median = 2.3 pA; n = 10) and in RyR3 -/- CA1 pyramidal neurons (mean ± SEM: 9.2 ± 3.4 pA; median = 4.7 pA; n = 7) was similar (P = 0.07, Mann-Whitney test). At steady state (15 min) the amplitude of sI_AHP_ was similar in RyR3 +/+ neurons (mean ± SEM: 45.2 ± 8.1 pA; median = 37.7 pA; n = 10) and in RyR3 -/- (mean ± SEM: 30.5 ± 6.1 pA; median = 27.1 pA; n = 7) (P = 0.2). This shows that a clear potentiation was observed for both RyR3 +/+ (P = 0.0006, paired t-test) and RyR3 -/- (P = 0.006, paired t-test) neurons, because the sI_AHP_ amplitude at steady state was clearly larger when compared with sI_AHP_ amplitude at the beginning of the recording. (**D)** Time course of relative amplitude increase of sI_AHP_ during the first 15 min of the recording, with current amplitudes normalised to the starting current measured at 0 min. The sI_AHP_ potentiation was overall larger in RyR3 +/+ than in RyR3 -/- CA1 pyramidal cells (two-way ANOVA with Bonferroni’s test, P < 0.001). (**E)** The time constant (″) of sI_AHP_ potentiation was obtained by fitting a mono-exponential function to the sI_AHP_ amplitude during the run-up phase of each individual experiment. The time constant of potentiation was faster in RyR3 -/- (mean ± SEM: 3.1 ± 0.7 min; median = 3.2 min; n = 7) compared to RyR3 +/+ CA1 neurons (mean ± SEM: 6.7 ± 1.1 min; median = 5.6 min; n = 10) (P = 0.043).

However, the sI_AHP_ amplitude at steady state was consistently and significantly larger than the sI_AHP_ amplitude at the beginning of recording in both RyR3 +/+ (P = 0.0006, n = 10, paired t-test) and in RyR3 $/$ CA1 pyramidal neurons (P = 0.006, n = 7, paired t-test) (Fig 6C), revealing that some activity-dependent potentiation of the current occurred both in the presence and in the absence of RyR3. Interestingly, the relative sI_AHP_ potentiation was more pronounced in RyR3 +/+ CA1 pyramidal neurons (two-way ANOVA, P < 0.001) (Fig 6D). This was further corroborated by comparing the ratio of the sI_AHP_ amplitude measured at 0 and 15 minutes, which was lower in RyR3 $/$ (5.0 ± 1.0, n = 7) than in RyR3 +/+ (43.5 ± 19.1, n = 10) neurons (P = 0.02, Mann-Whitney test) (Fig 7G). Additionally, the time constant of the sI_AHP_ potentiation was lower in RyR3 $/$ than in RyR3 +/+ CA1 neurons (P = 0.043) (Fig 6E). These results suggest that lack of RyR3 leads to a faster and reduced activity-dependent potentiation of sI_AHP_.

**Fig 7.**
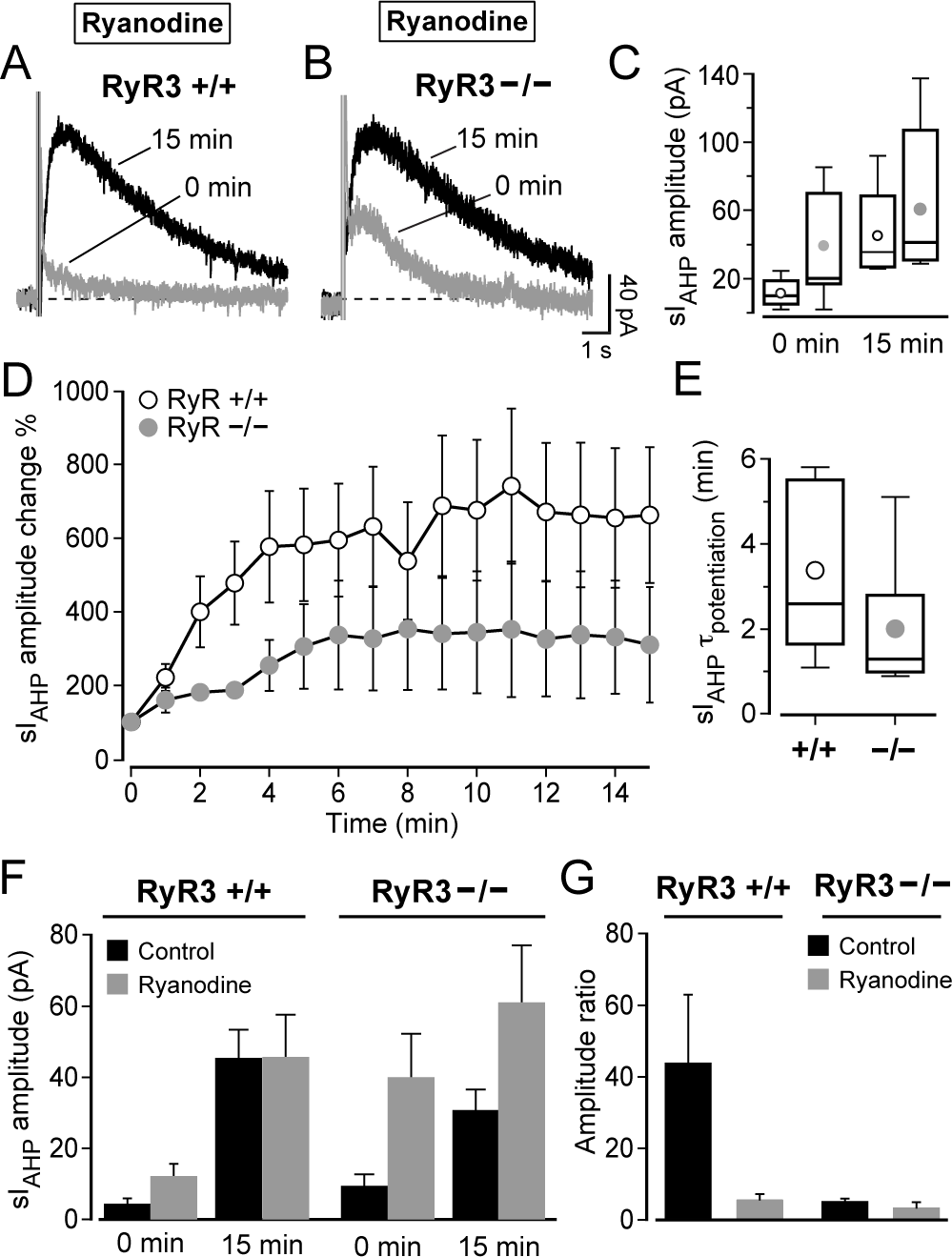
Type 3 ryanodine receptor (RyR3) is a main player in the CICR that mediates the activity-dependent potentiation of sI_AHP_ in mouse CA1 pyramidal neurons. (A, B) Superimposed traces of the sI_AHP_ recorded at 0 and 15 minutes from RyR3 +/+ (A) and RyR3 -/- (B) CA1 pyramidal neurons in the presence of 10 µM ryanodine from the outset of the recordings. (C) Summary box-and-whiskers plot of the sI_AHP_ amplitudes recorded at 0 min and at 15 min in the presence of ryanodine (10 µM). At 0 min the sI_AHP_ amplitudes in RyR3 +/+ CA1 neurons (mean ± SEM: 12.1 ± 3.7 pA; median = 10.4 pA; n = 5) and in RyR3 -/- CA1 pyramidal neurons (mean ± SEM: 39.9 ± 12.4 pA; median = 21.2 pA; n = 7) were not significantly different (P = 0.06, Mann-Whitney test). At 15 min the current in RyR3 +/+ (mean ± SEM: 45.6 ± 12.1 pA; median = 36.0 pA; n = 5) and in RyR3 -/- (mean ± SEM: 61.0 ± 16.3 pA; median = 41.6 pA; n = 7) was similar (P = 0.4, Mann- Whitney test). In the presence of ryanodine (10 µM) from the onset, RyR3 +/+ cells had a larger sI_AHP_ at 15 minutes than at 0 minute (P = 0.047, n = 5, t-test with Welch correction), while RyR3 -/- neurons had similar sI_AHP_ amplitudes at 0 and 15 minutes (P = 0.3, n = 7, Mann-Whitney test). (D) Time course of relative increase of sI_AHP_ during the first 15 min of the recording in the presence of ryanodine (10 µM), with current amplitudes normalized to the starting current measured at 0 min. The sI_AHP_ potentiation was overall similar in RyR3 +/+ and RyR3 -/- CA1 cells (P = 0.17, two-way repeated measures ANOVA). (E) The time constant (″) of sI_AHP_ potentiation in RyR3 +/+ CA1 neurons (mean ± SEM: 3.4 ± 0.9 min; median = 2.6 min; n = 5) and in RyR3 -/- (mean ± SEM: 2.0 ± 0.6 min; median = 1.3 min; n = 7) was similar (P = 0.2). (F) Summary bar chart comparing the sI_AHP_ amplitude from RyR3 +/+ and RyR3 -/- mice in the presence and absence of ryanodine. Ryanodine increased the initial amplitude of sI_AHP_ in RyR3 +/+ (P = 0.02, Mann-Whitney test) and RyR3 -/- CA1 neurons (P = 0.045, t-test with Welch correction), but it did not affect the current measured at 15 minutes in either RyR3 +/+ (P > 0.9) or RyR3 -/- CA1 neurons (P = 0.1, Mann-Whitney test). (G) Summary bar chart comparing ratios of the sI_AHP_ amplitude measured at 0 and 15 min from RyR3 +/+ and RyR3 -/- mice with or without ryanodine. Ryanodine did not significantly decrease the sI_AHP_ ratio in RyR3 -/- CA1 neurons, and did not affect the ratio in RyR3 +/+ CA1 neurons, in spite of an apparent trend (one-way ANOVA-Kruskal-Wallis test with Dunn’s multiple comparison test, P > 0.05 for both comparisons). In the presence of ryanodine from the onset, the sI_AHP_ ratio was not significantly different between RyR3 +/+ CA1 cells (5.5 ± 1.8, n = 5) and RyR3 -/- CA1 neurons (3.2 ± 1.8, n = 7) (one-way ANOVA-Kruskal-Wallis test with Dunn’s multiple comparison test, P > 0.05 for this comparison).

### CICR underlying the activity-dependent potentiation of sI_AHP_ is mostly dependent on RyR3 in mouse CA1 pyramidal neurons

Our results obtained from rat CA1 pyramidal neurons showed that inhibition of CICR by ryanodine and CPA application substantially reduced the activity-dependent potentiation of sI_AHP_ (Fig 3). If RyR3 is the molecular determinant for the activity-dependent potentiation of the sI_AHP_, then potentiation in RyR3 $/$ mice should not be affected by ryanodine.

In the presence of ryanodine (10 µM) from the start of the recording, in RyR3 +/+ CA1 neurons the sI_AHP_ amplitude significantly increased from an initial value of 12.1 ± 3.7 pA to 45.6 ± 12.1 pA, when the sI_AHP_ was fully potentiated (n = 5, P = 0.047, t-test with Welch correction) (Fig 7A, C and F). However, the sI_AHP_ amplitude in RyR3 $/$ CA1 neurons recorded at the beginning (39.9 ± 12.4 pA) was not significantly different from the amplitude recorded at the end of the run-up phase (61.0 ± 16.3 pA; n = 7, P = 0.3, Mann-Whitney test) (Fig 7B, C and F). Indeed, the change of sI_AHP_ amplitude throughout the run-up phase was comparable between RyR3 +/+ and RyR3 $/$ CA1 neurons (P = 0.17, two-way repeated measures ANOVA) (Fig 7D). Also, the time for the sI_AHP_ to reach full potentiation was not different between RyR3 +/+ and RyR3 $/$ CA1 cells (P = 0.2) (Fig 7E). This observed similarity in run-up in the presence of ryanodine is also seen when comparing the ratio of the sI_AHP_ amplitude recorded at 0 minutes and 15 minute, which was not significantly different between RyR3 +/+ and RyR3 $/$ CA1 neurons (one-way ANOVA-Kruskal-Wallis test with Dunn’s multiple comparison test, P > 0.05) (Fig 7G). The similarity of the relative extent of run-up (Fig 7D and E) and of the amplitude ratio (Fig 7G) indicates that no difference in sI_AHP_ potentiation between knock-out and wild-type animals was observed when ryanodine was applied from the beginning of the recordings.

When comparing the sI_AHP_ amplitudes recorded without ryanodine (Fig 6A-C and Fig 7F) with those where ryanodine was present from the beginning of the recording (Fig 7A-C and F) in RyR3 +/+ and RyR3 $/$ CA1 neurons, it becomes evident that the absence of RyR3 leads to a reduction in the activity-dependent potentiation of sI_AHP_ by increasing its initial amplitude. In the presence of ryanodine, the initial (0 min) sI_AHP_ amplitude was not quite significantly different between RyR3 +/+ (12.1 ± 3.7 pA) and RyR3 $/$ (39.9 ± 12.4 pA) CA1 neurons, in spite of a tendency towards larger currents in RyR3 $/$ cells, (P = 0.06, t-test with Welch correction) (Fig 7C and F). However, the initial sI_AHP_ amplitude (0 min) was significantly increased in the presence of ryanodine in RyR3 +/+ CA1 neurons (P = 0.02, Mann-Whitney test) and, to an even larger extent, in RyR3 $/$ (P = 0.045, t-test with Welch correction) (Fig 7F). Conversely, the sI_AHP_ amplitude measured at the end of the run-up phase (15 min) was not different between RyR3 +/+ and RyR3 $/$ CA1 cells (P = 0.4, Mann-Whitney test; Fig 7A-C and F), and was similar with and without ryanodine applied from the beginning of the recording in RyR3 +/+ CA1 neurons (P > 0.9) and in RyR3 $/$ CA1 neurons (P = 0.1, Mann-Whitney test) (Fig 7F). The increase in initial current amplitude, caused by ryanodine and particularly prominent in the absence of RyR3, combined with no change in the current amplitude at the end of the run-up phase results in the reduction in potentiation observed when reducing CICR by ryanodine, and especially pronounced in RyR3 $/$ CA1 neurons. Thus, overall inhibition of CICR by ryanodine had an effect on the starting but not final sI_AHP_ amplitudes both in the presence and absence of RyR3. Taken altogether, the data show that inhibition of CICR by ryanodine removed the differences in the extent and rate of potentiation of sI_AHP_ between RyR3 +/+ and RyR3 $/$ CA1 neurons, suggesting that type 3 ryanodine receptors play a specific role in mediating the activity-dependent potentiation of this current.

## Discussion

This study shows that ryanodine receptor-mediated calcium-induced calcium release contributes both to the amplitude of the sI_AHP_ at steady state and its activity-dependent potentiation in rat and mouse hippocampal pyramidal neurons. In particular, analysis of CA1 pyramidal neurons lacking type 3 ryanodine receptors (RyR3) has revealed that RyR3 plays an essential and specific role in shaping the activity-dependent potentiation of the sI_AHP_. RyR-mediated CICR contributes to the generation of currents underlying long-lasting afterhyperpolarisations in a variety of neurons, including guinea pig dorsal vagal nucleus neurons [20], sympathetic neurons [45], superior cervical ganglion neurons [46], nodose ganglion neurons [47], afterhyperpolarising (AH) myenteric plexus neurons in guinea pig ileum [48] and duodenum [49], rabbit otic ganglion neurons [50], and layer II-III sensorimotor neocortical completely adapting neurons [19]. In these studies ryanodine, applied at concentrations of 5-20 µM led to reductions in the amplitude of the sAHP and the corresponding sI_AHP_ ranging from ∼50% [19, 45, 48] to complete inhibition [20, 47, 49].

The contribution of RyR-mediated CICR to the generation of the sAHP or the sI_AHP_ in hippocampal neurons is more controversial. In CA3 pyramidal neurons from organotypic slice cultures sI_AHP_ was inhibited by ∼50% upon application of ryanodine [10], which is comparable to the ∼30% reduction observed in CA1-CA3 cultured hippocampal neurons [51], suggesting that CICR plays a role in the activation of sI_AHP_. In CA1 pyramidal neurons, our study reports a reduction of the sI_AHP_ of 23.3 ± 3.7% (Fig 2E) from rat and 39.0 ± 6.9 % (Fig 5I) from mouse hippocampal slices in response to the application of ryanodine once the current had reached its steady-state amplitude.

Therefore our results are in good agreement with those in CA3 and cultured hippocampal neurons [10, 51], and with other studies on CA1 pyramidal neurons in slices [14, 28, 29, 52], but differ from those of Zhang et al. (1995) and Torres et al. (1996), who failed to observe any effect of ryanodine on sI_AHP_ and sAHP in CA1 neurons, although Torres et al. (1996) reported a reduction in sAHP in response to other CICR inhibitors (dantrolene and ruthenium red).

The concentration of ryanodine used in our study is in a range that changes the conductance properties of RyR receptors, keeping them in a subconducting state with high open probability, thereby depleting the ryanodine sensitive Ca^2+^ stores and preventing further Ca^2+^ release [44, 53]. In a physiological setting this has been shown to reduce both the amplitude and duration of evoked Ca^2+^ transients and sAHP in myenteric neurons [48], to decrease the amplitude of action potential-evoked Ca^2+^ transients [27] and inhibit caffeine-induced Ca^2+^ signals [26] in CA1 pyramidal neurons. The time course of the effect of ryanodine on the fully developed sI_AHP_ showed that a clear sI_AHP_ inhibition was achieved in 10-20 min in our experiments, similar to what was observed in cultured hippocampal neurons [51] and vagal motorneurons [20].

The contribution of CICR to the generation of sI_AHP_ in CA1 neurons is further supported by our finding that caffeine, which activates ryanodine receptors causing calcium release [36, 54], enhances the sI_AHP_ amplitude. This finding is in line with similar observations in CA3 [55] and CA1 pyramidal neurons [30], but differs from the lack of effect of caffeine reported by Zhang et al. (1995). In AH myenteric neurons, application of caffeine (5 mM) was usually accompanied by a transient increase in sI_AHP_, followed by a decrease of the current [49]. This is similar to what we observed in CA1 pyramidal neurons upon application of caffeine at a lower concentration (0.5 mM; Fig 1A-E). Furthermore, our finding suggests that the observed decrease in sI_AHP_ amplitude results from the inhibition of phosphodiesterases by caffeine [40]. This leads to a slow but significant increase in cAMP and activation of protein kinase A (PKA), which is known to suppress the sI_AHP_ [42]. This is further supported by our evidence that inhibition of PKA by Rp-cAMPS prevents the caffeine-mediated reduction of sI_AHP_ (Fig 1F-J). Consequently, caffeine exerts a purely enhancing effect on the current under PKA inhibition, as expected upon increase in intracellular Ca^2+^ levels.

Our observations further suggest that PKA activation, as a consequence of phosphodiesterase inhibition, might underlie the inhibitory effect of caffeine on the sI_AHP_ in different types of neurons, as reported in other studies [20, 49].

Low concentrations of caffeine increase action potential-induced Ca^2+^ transients in CA1 pyramidal neurons, and this effect is dependent on facilitation of CICR and independent from activation of the cAMP-PKA pathway [27], similar to what we have shown for the sI_AHP_ modulation by caffeine. Taken together, these findings suggest that caffeine might enhance sAHP and spike frequency adaptation by facilitating CICR and increasing action potential-induced Ca^2+^ transients under physiological conditions in CA1 neurons.

Taken altogether, the effects of pharmacological modulators of CICR in our study suggest that ryanodine receptor-mediated CICR contributes to the generation of sI_AHP_ and to its amplitude at steady state. To further address the role of specific RyR subtypes in the generation of sI_AHP_, we have analysed the current in CA1 pyramidal neurons from mice lacking type 3 ryanodine receptors (RyR3 $/$). The salient features of sI_AHP_ (amplitude, charge transfer, time course of decay) were similar in the presence or absence of RyR3, despite a reduction in current density that was linked to a concomitant increase in membrane capacity in RyR3 deficient mice. Our finding suggests that the absence of RyR3 does not significantly impair the generation of sI_AHP_ and RyR3 does not significantly contribute to the absolute size of this current at the steady state. This finding contrasts with a report that intracellular application of anti-RyR3 antibodies caused a 70% reduction in sI_AHP_ and occluded the effect of ryanodine on this current in mouse CA1 pyramidal neurons [31]. In our experiments, ryanodine reduced the sI_AHP_ amplitude similarly in neurons from RyR3 +/+ (39.0 ± 6.9 %) and RyR3 $/$ (36.9 ± 9.4%) mice (Fig 5). Possible reasons for this discrepancy might be due to the different experimental conditions, such as the voltage protocols used to elicit sI_AHP_; in particular, the 1 s-long depolarising voltage steps used by Van de Vrede et al. (2007) have been shown to cause “overcharging” of ryanodine-sensitive stores in mouse CA1 pyramidal neurons [25], which might have led to an over-estimation of the contribution of RyR-mediated calcium release to the sI_AHP_ under those experimental conditions when compared to more physiological depolarising stimuli to elicit this current. The other experimental difference is the use of a specific knock-out missing the RyR3 receptor instead of the intracellular application of antibody [31]. In the absence of an independent validation of the specific blocking effect of the anti-RyR3 antibody on RyR3 channel activity [56], it cannot be excluded that the pronounced effect of the antibody on sI_AHP_[31] might be due partly to off-target effects. In the case of the RyR3 knock-out mice, it is worth noticing that, even when multiple subtypes of ryanodine receptors are expressed in the same cell type (RyR1, RyR2 and RyR3 in the case of CA1 pyramidal neurons) [22–24], available experimental evidence supports the formation of homomeric channels [57], and knockout of the RyR3 gene does not affect the expression level of the RyR1 and RyR2 isoforms in the brain [58, 59]. Thus, the lack of significant changes in the steady state size of the sI_AHP_ and its modulation by ryanodine in RyR3 $/$ compared to RyR3 (+/+) (Fig 4 and 5) is unlikely to be due to a compensatory up-regulation of RyR1 or RyR2 expression in CA1 neurons.

Repeated activation of CA1 pyramidal neurons with depolarising current pulses [15] or action potential firing at gamma-related (50 Hz) frequencies [16] causes a long-lasting, gradual decrease in membrane excitability associated with an increase in spike frequency adaptation and a potentiation of the sAHP. Additionally, voltage-clamp analysis has revealed a concomitant activity-dependent potentiation of sI_AHP_ in rat CA1 pyramidal neurons, critically dependent on the activation of L-type voltage-gated Ca^2+^ channels and ryanodine-sensitive CICR [14]. Our results support the findings of Borde et al. (2000) and extend them to mouse CA1 pyramidal neurons. Most importantly, a detailed analysis of the sI_AHP_ potentiation in CA1 neurons lacking RyR3 receptors has revealed that, in the absence of RyR3, the activity-dependent potentiation of sI_AHP_ is reduced and its time course is faster than in CA1 neurons expressing RyR3 (Fig 6). The differences in the extent and rate of potentiation of sI_AHP_ between RyR3 +/+ and RyR3 $/$ CA1 neurons are removed by overall inhibition of CICR by ryanodine (Fig 7). Our findings suggest that type 3 ryanodine receptors are not essential for the generation and maintenance of sI_AHP_, but play a specific and distinctive role in shaping the extent and time course of the activity-dependent potentiation of this current. Our data, however, do not provide any direct indication as to how ryanodine-sensitive CICR, and in particular RyR3, contribute to the activity-dependent potentiation of the sI_AHP_. A potential mechanism might be linked to a distinctive property of RyR3 compared to other ryanodine receptors, namely their low sensitivity to inactivation at high calcium concentration [25, 60]. The resulting sustained RyR3 activity at elevated Ca^2+^ concentrations would make the RyR3 channels particularly suitable to provide a more sustained calcium release efflux from the endoplasmic reticulum when stimulated by increasing concentration of Ca^2+^ released by other calcium release channels, or by calcium channels on the plasma membrane [58]. This could also explain the shorter time course of sI_AHP_ potentiation observed in CA1 neurons in the absence of RyR3 (Fig 6D and E).

A second mechanism might result from the interaction between VGCCs in the plasma membrane and RyR-dependent calcium release. In CA1 pyramidal neurons L-type VGCCs are functionally coupled to RyR3, providing the Ca^2+^ influx necessary for RyR3-mediated CICR [61], and to the activation of sI_AHP_ [7, 9]. Ca^2+^ -dependent inactivation is a negative feedback mechanism whereby Ca^2+^ ions restrict their own entry into the cell by one of the main routes of Ca^2+^ influx, the VGCCs [62]. In thalamo-cortical relay neurons RyR-dependent CICR enhances the Ca^2+^-dependent inactivation of L-type VGCCs [63]. If a similar interplay is present between L-type Ca^2+^ channels and RyR-mediated CICR in CA1 neurons, this might potentially explain the increase in the amplitude of sI_AHP_ at the start of the recordings observed in the presence of ryanodine, especially in RyR3 deficient neurons (Fig 7F). A reduction of CICR in the absence of RyR3 receptors would lead to a reduced Ca^2+^-dependent inactivation of the L-type Ca^2+^ current, eliciting in turn a larger initial sI_AHP_.

A third potential mechanism might result from a different mode of activation of RyR channels, namely store-overload-induced Ca^2+^ release (SOICR) [64]. The repetitive stimulation leading to Ca^2+^ influx and underlying the activity-dependent potentiation of sI_AHP_ might increase Ca^2+^ uptake by the endoplasmic reticulum, resulting in Ca^2+^ overload and subsequently activation of RyRs by luminal Ca^2+^ and SOICR, in analogy to what has been observed in cardiac cells [65]. SOICR has primarily been shown to be mediated by RyR2, with RyR1 having a lower sensitivity, in cardiac and skeletal muscle cells, as well as in HEK293 cells expressing recombinant RyR1 and RyR2 [66], but has not been shown in neurons or linked to RyR3 function. However, a store Ca^2+^ sensing gate structure has been identified in RyR2 and is conserved in all types of RyRs [67]. In view of this structural similarity, it is tempting to speculate that RyR3 might mediate SOICR in response to repetitive activity in CA1 neurons, and this might specifically contribute to the activity-dependent potentiation of sI_AHP_.

The modulation of activity-dependent potentiation of sI_AHP_ by RyR3-mediated CICR shown in this study is likely to contribute to plasticity of intrinsic neuronal excitability and act postsynaptically to control the flow of synaptic signals [13], regulate the threshold for induction of long-term potentiation in hippocampal neurons [68], and play a critical role in learning and memory [69–71].

## Acknowledgments

We thank Nicole Dalgleish for help with preliminary data analysis.

## Author Contributions

**Conceptualization:** Paola Pedarzani, Martin Stocker

**Formal analysis:** Petra Ludwig, Paola Pedarzani, Angelo Tedoldi

**Funding acquisition:** Paola Pedarzani, Martin Stocker, Hiroshi Takeshima

**Methodology:** Paola Pedarzani, Angelo Tedoldi

**Investigation:** Gianluca Fulgenzi, Petra Ludwig, Martin Stocker, Angelo Tedoldi

**Project administration:** Paola Pedarzani, Martin Stocker

**Resources:** Hiroshi Takeshima

**Supervision:** Paola Pedarzani, Martin Stocker

**Validation:** Paola Pedarzani

**Visualisation:** Petra Ludwig, Paola Pedarzani, Martin Stocker, Angelo Tedoldi

**Writing – original draft:** Paola Pedarzani, Martin Stocker

**Writing – review & editing:** Gianluca Fulgenzi, Petra Ludwig, Paola Pedarzani, Martin Stocker, Hiroshi Takeshima, Angelo Tedoldi

## References

1. Alger BE, Nicoll RA. Epileptiform burst afterhyperolarization: calcium-dependent potassium potential in hippocampal CA1 pyramidal cells. Science. 1980; 210(4474): 1122–1124. doi: 10.1126/science.7444438 PMID: 7444438.

2. Hotson JR, Prince DA. A calcium-activated hyperpolarization follows repetitive firing in hippocampal neurons. J Neurophysiol. 1980; 43(2): 409–419. doi: 10.1152/jn.1980.43.2.409 PMID: 6247461.

3. Madison DV, Nicoll RA. Noradrenaline blocks accommodation of pyramidal cell discharge in the hippocampus. Nature. 1982; 299(5884): 636-638. doi: 10.1038/299636a0 PMID: 6289127.

4. Madison DV, Nicoll RA. Control of the repetitive discharge of rat CA 1 pyramidal neurones in vitro. J Physiol (Lond). 1984; 354: 319–331. doi: 10.1113/jphysiol.1984.sp015378 PMID: 6434729.

5. Lancaster B, Adams PR. Calcium-dependent current generating the afterhyperpolarization of hippocampal neurons. J Neurophysiol. 1986; 55(6): 1268–1282. doi: 10.1152/jn.1986.55.6.1268 PMID: 2426421.

6. Sah P. Ca^2+^-activated K^+^ currents in neurones: types, physiological roles and modulation. Trends Neurosci. 1996; 19(4): 150–154. doi: 10.1016/s0166-2236(96)80026-9 PMID: 8658599.

7. Lima PA, Marrion NV. Mechanisms underlying activation of the slow AHP in rat hippocampal neurons. Brain Res. 2007; 1150: 74–82. doi: 10.1016/j.brainres.2007.02.067 PMID: 17395164.

8. Moyer JR, Jr., Thompson LT, Black JP, Disterhoft JF. Nimodipine increases excitability of rabbit CA1 pyramidal neurons in an age- and concentration-dependent manner. J Neurophysiol. 1992; 68(6): 2100–2109. doi: 10.1152/jn.1992.68.6.2100 PMID: 1491260.

9. Rascol O, Potier B, Lamour Y, Dutar P. Effects of calcium channel agonist and antagonists on calcium-dependent events in CA1 hippocampal neurons. Fundam Clin Pharmacol. 1991; 5(4): 299–317. doi: 10.1111/j.1472-8206.1991.tb00725.x PMID: 1717356.

10. Tanabe M, Gähwiler BH, Gerber U. L-Type Ca^2+^ channels mediate the slow Ca^2+^-dependent afterhyperpolarization current in rat CA3 pyramidal cells in vitro. J Neurophysiol. 1998; 80(5): 2268–2273. doi: 10.1152/jn.1998.80.5.2268 PMID: 9819242.

11. Gamelli AE, McKinney BC, White JA, Murphy GG. Deletion of the L-type calcium channel Ca_v_1.3 but not Ca_v_1.2 results in a diminished sAHP in mouse CA1 pyramidal neurons. Hippocampus. 2011; 21(2): 133–141. doi: 10.1002/hipo.20728 PMID: 20014384.

12. Jahromi BS, Zhang L, Carlen PL, Pennefather P. Differential time-course of slow afterhyperpolarizations and associated Ca^2+^ transients in rat CA1 pyramidal neurons: further dissociation by Ca^2+^ buffer. Neuroscience. 1999; 88(3): 719–726. doi: 10.1016/s0306-4522(98)00203-6 PMID: 10363812.

13. Borde M, Bonansco C, Buño W. The activity-dependent potentiation of the slow Ca^2+^-activated K^+^ current regulates synaptic efficacy in rat CA1 pyramidal neurons. Pflugers Arch. 1999; 437(2): 261–266. doi: 10.1007/s004240050778 PMID: 9929568.

14. Borde M, Bonansco C, Fernandez de Sevilla D, Le Ray D, Buño W. Voltage-clamp analysis of the potentiation of the slow Ca^2+^-activated K^+^ current in hippocampal pyramidal neurons. Hippocampus. 2000; 10(2): 198–206. doi: 10.1002/(sici)1098-1063(2000)10:2<198::aid- hipo9>3.0.co;2-f PMID: 10791842.

15. Borde M, Cazalets JR, Buño W. Activity-dependent response depression in rat hippocampal CA1 pyramidal neurons in vitro. J Neurophysiol. 1995; 74(4): 1714–1729. doi: 10.1152/jn.1995.74.4.1714 PMID: 8989407.

16. Kaczorowski CC. Bidirectional pattern-specific plasticity of the slow afterhyperpolarization in rats: role for high-voltage activated Ca^2+^ channels and I. Eur J Neurosci. 2011; 34(11): 1756–1765. doi: 10.1111/j.1460-9568.2011.07899.x PMID: 22098477.

17. Zhang L, Pennefather P, Velumian A, Tymianski M, Charlton M, Carlen PL. Potentiation of a slow Ca^2+^-dependent K^+^ current by intracellular Ca^2+^ chelators in hippocampal CA1 neurons of rat brain slices. J Neurophysiol. 1995; 74(6): 2225–2241. doi: 10.1152/jn.1995.74.6.2225 PMID: 8747186.

18. Akita T, Kuba K. Functional triads consisting of ryanodine receptors, Ca^2+^ channels, and Ca^2+^- activated K^+^ channels in bullfrog sympathetic neurons. Plastic modulation of action potential. J Gen Physiol. 2000; 116(5): 697–720. doi: 10.1085/jgp.116.5.697 PMID: 11055998.

19. Pineda JC, Galarraga E, Foehring RC. Different Ca2+ source for slow AHP in completely adapting and repetitive firing pyramidal neurons. Neuroreport. 1999; 10(9): 1951–1956. doi: 10.1097/00001756-199906230-00029 PMID: 10501539.

20. Sah P, McLachlan EM. Ca^2+^-activated K^+^ currents underlying the afterhyperpolarization in guinea pig vagal neurons: a role for Ca^2+^-activated Ca^2+^ release. Neuron. 1991; 7(2): 257–264. doi: 10.1016/0896-6273(91)90264-z PMID: 1873029.

21. Bardo S, Cavazzini MG, Emptage N. The role of the endoplasmic reticulum Ca^2+^ store in the plasticity of central neurons. Trends Pharmacol Sci. 2006; 27(2): 78–84. doi: 10.1016/j.tips.2005.12.008 PMID: 16412523.

22. Furuichi T, Furutama D, Hakamata Y, Nakai J, Takeshima H, Mikoshiba K. Multiple types of ryanodine receptor/Ca^2+^ release channels are differentially expressed in rabbit brain. J Neurosci. 1994; 14(8): 4794–4805. doi: 10.1523/jneurosci.14-08-04794.1994 PMID: 8046450.

23. Giannini G, Conti A, Mammarella S, Scrobogna M, Sorrentino V. The ryanodine receptor/calcium channel genes are widely and differentially expressed in murine brain and peripheral tissues. J Cell Biol. 1995; 128(5): 893–904. doi: 10.1083/jcb.128.5.893 PMID: 7876312.

24. Mori F, Fukaya M, Abe H, Wakabayashi K, Watanabe M. Developmental changes in expression of the three ryanodine receptor mRNAs in the mouse brain. Neurosci Lett. 2000; 285(1): 57–60. doi: 10.1016/s0304-3940(00)01046-6 PMID: 10788707.

25. Chen-Engerer H-J, Hartmann J, Karl RM, Yang J, Feske S, Konnerth A. Two types of functionally distinct Ca^2+^ stores in hippocampal neurons. Nat Commun. 2019; 10(1): 3223–3223. doi: 10.1038/s41467-019-11207-8 PMID: 31324793.

26. Garaschuk O, Yaari Y, Konnerth A. Release and sequestration of calcium by ryanodine- sensitive stores in rat hippocampal neurones. J Physiol (Lond). 1997; 502 (Pt 1)(Pt 1): 13–30. doi: 10.1111/j.1469-7793.1997.013bl.x PMID: 9234194.

27. Sandler VM, Barbara JG. Calcium-induced calcium release contributes to action potential- evoked calcium transients in hippocampal CA1 pyramidal neurons. J Neurosci. 1999; 19(11): 4325–4336. doi: 10.1523/jneurosci.19-11-04325.1999 PMID: 10341236.

28. Gant JC, Sama MM, Landfield PW, Thibault O. Early and simultaneous emergence of multiple hippocampal biomarkers of aging is mediated by Ca^2+^-induced Ca^2+^ release. J Neurosci. 2006; 26(13): 3482–3490. doi: 10.1523/jneurosci.4171-05.2006 PMID: 16571755.

29. Kumar A, Foster TC. Enhanced long-term potentiation during aging is masked by processes involving intracellular calcium stores. J Neurophysiol. 2004; 91(6): 2437–2444. doi: 10.1152/jn.01148.2003 PMID: 14762159.

30. Torres GE, Arfken CL, Andrade R. 5-Hydroxytryptamine4 receptors reduce afterhyperpolarization in hippocampus by inhibiting calcium-induced calcium release. Mol Pharmacol. 1996; 50(5): 1316–1322. PMID: 8913363.

31. van de Vrede Y, Fossier P, Baux G, Joels M, Chameau P. Control of IsAHP in mouse hippocampus CA1 pyramidal neurons by RyR3-mediated calcium-induced calcium release. Pflugers Arch. 2007; 455(2): 297–308. doi: 10.1007/s00424-007-0277-4 PMID: 17562071.

32. Takeshima H, Ikemoto T, Nishi M, Nishiyama N, Shimuta M, Sugitani Y, Kuno J, Saito I, Saito H, Endo M, Iino M, Noda T. Generation and characterization of mutant mice lacking ryanodine receptor type 3. J Biol Chem. 1996; 271(33): 19649–19652. doi: 10.1074/jbc.271.33.19649 PMID: 8702664.

33. Blanton MG, Lo Turco JJ, Kriegstein AR. Whole cell recording from neurons in slices of reptilian and mammalian cerebral cortex. J Neurosci Methods. 1989; 30(3): 203–210. PMID: 2607782.

34. Rothman JS, Silver RA. NeuroMatic: An Integrated Open-Source Software Toolkit for Acquisition, Analysis and Simulation of Electrophysiological Data. Front Neuroinform. 2018; 12: 14–14. doi: 10.3389/fninf.2018.00014 PMID: 29670519.

35. Stocker M, Krause M, Pedarzani P. An apamin-sensitive Ca^2+^-activated K^+^ current in hippocampal pyramidal neurons. Proc Natl Acad Sci U S A. 1999; 96(8): 4662–4667. PMID: 10200319.

36. Thomas NL, Williams AJ. Pharmacology of ryanodine receptors and Ca^2+^-induced Ca^2+^ release. Wiley Interdisciplinary Reviews: Membrane and Transport Signaling. 2012; 1(4): 383–397. doi: 10.1002/wmts.34.

37. McPherson PS, Kim YK, Valdivia H, Knudson CM, Takekura H, Franzini-Armstrong C, Coronado R, Campbell KP. The brain ryanodine receptor: a caffeine-sensitive calcium release channel. Neuron. 1991; 7(1): 17–25. doi: 10.1016/0896-6273(91)90070-g PMID: 1648939.

38. Sitsapesan R, Williams AJ. Mechanisms of caffeine activation of single calcium-release channels of sheep cardiac sarcoplasmic reticulum. J Physiol (Lond). 1990; 423: 425–439. doi: 10.1113/jphysiol.1990.sp018031 PMID: 2167363.

39. Butcher RW, Sutherland EW. Adenosine 3’,5’-phosphate in biological materials. I. Purification and properties of cyclic 3’,5’-nucleotide phosphodiesterase and use of this enzyme to characterize adenosine 3’,5’-phosphate in human urine. J Biol Chem. 1962; 237: 1244–1250. PMID: 13875173.

40. Sawynok J, Yaksh TL. Caffeine as an analgesic adjuvant: a review of pharmacology and mechanisms of action. Pharmacol Rev. 1993; 45(1): 43–85. PMID: 8475169.

41. Pedarzani P, Krause M, Haug T, Storm JF, Stuhmer W. Modulation of the Ca^2+^-activated K^+^ current sIAHP by a phosphatase-kinase balance under basal conditions in rat CA1 pyramidal neurons. J Neurophysiol. 1998; 79(6): 3252–3256. doi: 10.1152/jn.1998.79.6.3252 PMID: 9636123.

42. Pedarzani P, Storm JF. PKA mediates the effects of monoamine transmitters on the K^+^ current underlying the slow spike frequency adaptation in hippocampal neurons. Neuron. 1993; 11(6): 1023–1035. doi: 10.1016/0896-6273(93)90216-e PMID: 8274274.

43. Botelho LH, Webster LC, Rothermel JD, Baraniak J, Stec WJ. Inhibition of cAMP-dependent protein kinase by adenosine cyclic 3’-, 5’-phosphorodithioate, a second cAMP antagonist. J Biol Chem. 1988; 263(11): 5301–5305. PMID: 2833504.

44. Rousseau E, Smith JS, Meissner G. Ryanodine modifies conductance and gating behavior of single Ca^2+^ release channel. Am J Physiol. 1987; 253(3 Pt 1): C364-368. doi: 10.1152/ajpcell.1987.253.3.C364 PMID: 2443015.

45. Jobling P, McLachlan EM, Sah P. Calcium induced calcium release is involved in the afterhyperpolarization in one class of guinea pig sympathetic neurone. J Auton Nerv Syst. 1993; 42(3): 251–257. doi: 10.1016/0165-1838(93)90370-a PMID: 8459099.

46. Davies PJ, Ireland DR, McLachlan EM. Sources of Ca^2+^ for different Ca^2+^-activated K^+^ conductances in neurones of the rat superior cervical ganglion. J Physiol (Lond). 1996; 495 (Pt 2): 353–366. doi: 10.1113/jphysiol.1996.sp021599 PMID: 8887749.

47. Cordoba-Rodriguez R, Moore KA, Kao JP, Weinreich D. Calcium regulation of a slow post- spike hyperpolarization in vagal afferent neurons. Proc Natl Acad Sci U S A. 1999; 96(14): 7650–7657. doi: 10.1073/pnas.96.14.7650 PMID: 10393875

48. Hillsley K, Kenyon JL, Smith TK. Ryanodine-sensitive stores regulate the excitability of AH neurons in the myenteric plexus of guinea-pig ileum. J Neurophysiol. 2000; 84(6): 2777–2785. doi: 10.1152/jn.2000.84.6.2777 PMID: 11110808.

49. Vogalis F, Furness JB, Kunze WA. Afterhyperpolarization current in myenteric neurons of the guinea pig duodenum. J Neurophysiol. 2001; 85(5): 1941–1951. doi: 10.1152/jn.2001.85.5.1941 PMID: 11353011.

50. Yoshizaki K, Hoshino T, Sato M, Koyano H, Nohmi M, Hua SY, Kuba K. Ca^2+^-induced Ca^2+^ release and its activation in response to a single action potential in rabbit otic ganglion cells. J Physiol (Lond). 1995; 486 (Pt 1): 177–187. doi: 10.1113/jphysiol.1995.sp020801 PMID: 7562634.

51. Shah M, Haylett DG. Ca^2+^ channels involved in the generation of the slow afterhyperpolarization in cultured rat hippocampal pyramidal neurons. J Neurophysiol. 2000; 83(5): 2554–2561. doi: 10.1152/jn.2000.83.5.2554 PMID: 10805657.

52. Gant JC, Chen KC, Norris CM, Kadish I, Thibault O, Blalock EM, Porter NM, Landfield PW. Disrupting function of FK506-binding protein 1b/12.6 induces the Ca^2+^-dysregulation aging phenotype in hippocampal neurons. J Neurosci. 2011; 31(5): 1693–1703. doi: 10.1523/JNEUROSCI.4805-10.2011 PMID: 21289178.

53. Tinker A, Sutko JL, Ruest L, Deslongchamps P, Welch W, Airey JA, Gerzon K, Bidasee KR, Besch HR, Jr., Williams AJ. Electrophysiological effects of ryanodine derivatives on the sheep cardiac sarcoplasmic reticulum calcium-release channel. Biophys J. 1996; 70(5): 2110–2119. doi: 10.1016/S0006-3495(96)79777-1 PMID: 9172735.

54. Meissner G. The structural basis of ryanodine receptor ion channel function. J Gen Physiol. 2017; 149(12): 1065–1089. doi: 10.1085/jgp.201711878 PMID: 29122978.

55. Qin Z, Zhou X, Gomez-Smith M, Pandey NR, Lee KF, Lagace DC, Beique JC, Chen HH. LIM domain only 4 (LMO4) regulates calcium-induced calcium release and synaptic plasticity in the hippocampus. J Neurosci. 2012; 32(12): 4271–4283. doi: 10.1523/JNEUROSCI.6271-11.2012 PMID: 22442089.

56. Naylor J, Beech DJ. Extracellular Ion Channel Inhibitor Antibodies. The Open Drug Discovery Journal. 2009; 1(1): 36–42. doi: 10.2174/1877381800901010036.

57. Murayama T, Ogawa Y. Characterization of type 3 ryanodine receptor (RyR3) of sarcoplasmic reticulum from rabbit skeletal muscles. J Biol Chem. 1997; 272(38): 24030–24037. doi: 10.1074/jbc.272.38.24030 PMID: 9295356.

58. Balschun D, Wolfer DP, Bertocchini F, Barone V, Conti A, Zuschratter W, Missiaen L, Lipp HP, Frey JU, Sorrentino V. Deletion of the ryanodine receptor type 3 (RyR3) impairs forms of synaptic plasticity and spatial learning. EMBO J. 1999; 18(19): 5264–5273. doi: 10.1093/emboj/18.19.5264 PMID: 10508160.

59. Liu J, Supnet C, Sun S, Zhang H, Good L, Popugaeva E, Bezprozvanny I. The role of ryanodine receptor type 3 in a mouse model of Alzheimer disease. Channels (Austin). 2014; 8(3): 230–242. doi: 10.4161/chan.27471 PMID: 24476841.

60. Chen SR, Li X, Ebisawa K, Zhang L. Functional characterization of the recombinant type 3 Ca^2+^ release channel (ryanodine receptor) expressed in HEK293 cells. J Biol Chem. 1997; 272(39): 24234–24246. doi: 10.1074/jbc.272.39.24234 PMID: 9305876.

61. Huddleston AT, Tang W, Takeshima H, Hamilton SL, Klann E. Superoxide-induced potentiation in the hippocampus requires activation of ryanodine receptor type 3 and ERK. J Neurophysiol. 2008; 99(3): 1565–1571. doi: 10.1152/jn.00659.2007 PMID: 18199822.

62. Budde T, Meuth S, Pape HC. Calcium-dependent inactivation of neuronal calcium channels. Nat Rev Neurosci. 2002; 3(11): 873–883. doi: 10.1038/nrn959 PMID: 12415295.

63. Rankovic V, Ehling P, Coulon P, Landgraf P, Kreutz MR, Munsch T, Budde T. Intracellular Ca^2+^ release-dependent inactivation of Ca^2+^ currents in thalamocortical relay neurons. Eur J Neurosci. 2010; 31(3): 439–449. doi: 10.1111/j.1460-9568.2010.07081.x PMID: 20105233.

64. Jiang D, Xiao B, Yang D, Wang R, Choi P, Zhang L, Cheng H, Chen SR. RyR2 mutations linked to ventricular tachycardia and sudden death reduce the threshold for store-overload-induced Ca^2+^ release (SOICR). Proc Natl Acad Sci U S A. 2004; 101(35): 13062–13067. doi: 10.1073/pnas.0402388101 PMID: 15322274.

65. Priori SG, Chen SR. Inherited dysfunction of sarcoplasmic reticulum Ca^2+^ handling and arrhythmogenesis. Circ Res. 2011; 108(7): 871–883. doi: 10.1161/CIRCRESAHA.110.226845 PMID: 21454795.

66. Kong H, Wang R, Chen W, Zhang L, Chen K, Shimoni Y, Duff HJ, Chen SR. Skeletal and cardiac ryanodine receptors exhibit different responses to Ca^2+^ overload and luminal Ca^2+^. Biophys J. 2007; 92(8): 2757–2770. doi: 10.1529/biophysj.106.100545 PMID: 17259277.

67. Chen W, Wang R, Chen B, Zhong X, Kong H, Bai Y, Zhou Q, Xie C, Zhang J, Guo A, Tian X, Jones PP, O’Mara ML, Liu Y, Mi T, Zhang L, Bolstad J, Semeniuk L, Cheng H, Chen J, Tieleman DP, Gillis AM, Duff HJ, Fill M, Song LS, Chen SR. The ryanodine receptor store- sensing gate controls Ca^2+^ waves and Ca^2+^-triggered arrhythmias. Nat Med. 2014; 20(2): 184–192. doi: 10.1038/nm.3440 PMID: 24441828.

68. Sah P, Bekkers JM. Apical dendritic location of slow afterhyperpolarization current in hippocampal pyramidal neurons: implications for the integration of long-term potentiation. J Neurosci. 1996; 16(15): 4537–4542. PMID: 8764642.

69. Oh MM, Oliveira FA, Disterhoft JF. Learning and aging related changes in intrinsic neuronal excitability. Front Aging Neurosci. 2010; 2: 2. doi: 10.3389/neuro.24.002.2010 PMID: 20552042.

70. Saar D, Barkai E. Long-term modifications in intrinsic neuronal properties and rule learning in rats. Eur J Neurosci. 2003; 17(12): 2727–2734. doi: 10.1046/j.1460-9568.2003.02699.x PMID: 12823479.

71. Zhang W, Linden DJ. The other side of the engram: experience-driven changes in neuronal intrinsic excitability. Nat Rev Neurosci. 2003; 4(11): 885–900. doi: 10.1038/nrn1248 PMID: 14595400.

